# A Scribble-E-cadherin complex controls daughter cell patterning by multiple mechanisms

**DOI:** 10.1101/2021.04.15.440081

**Authors:** Anchi S. Chann, Ye Chen, Tanja Kinwel, Patrick O. Humbert, Sarah M. Russell

## Abstract

The fate of the two daughter cells is intimately connected to their positioning, which is in turn regulated by cell junction remodelling and orientation of the mitotic spindle. How multiple cues are integrated to dictate the ultimate patterning of daughters is not clear. Here, we identify novel mechanisms of regulation of daughter positioning in single MCF10A cells. The polarity protein, Scribble, links E-cadherin to NuMA and Arp2/3 signalling for sequential roles in daughter positioning. First Scribble transmits cues from E-cadherin localised in retraction fibres to control orientation of the mitotic spindle. Second, Scribble re-locates to the junction between the two daughters to allow a new E-cadherin-based-interface to form between them, influencing the width of the nascent daughter-daughter junction, generation of filopodia and subsequent cell patterning. Thus, E-cadherin and Scribble dynamically relocate to different intracellular sites during cell division to orient the mitotic spindle and control placement of the daughter cells after cell division.

## Introduction

The position of the two daughter cells following cell division is critical to many biological processes, from embryonic patterning and tissue growth to the extrusion of mutant cells^1, 2^. Daughter cell positioning is coordinated by a complex interplay of responses to intrinsic and extrinsic cues. These cues can include receptors at cell-cell contact sites and the extracellular matrix, and non-chemical influences such as cell geometry and tissue tension^3^. Cues are translated to daughter cell positioning through cell activities such as remodelling of junctions and orienting of the mitotic spindle^4^. An active question in the field is how these cues enable memory of the original position of the parent, and are transmitted to control position and fate of the daughter cells.

Spindle orientation provides a functional link between spatial context and fate of the progeny of a cell division in many contexts. For instance, spindle orientation influences the position, the size, and the fate of the two daughter cells of an epithelial division, impacting upon cell diversification and tissue homeostasis and morphogenesis^5–9^. Errors in the control of orientation of the mitotic spindle lead to developmental defects and cancer^5, 10^. Recent studies indicate that several attributes combine to influence spindle orientation and daughter cell positioning, including intrinsic polarity, the location of adhesions and constraints on the cell shape^11–22^. All these cues are transmitted to spindle orientation via positioning of the LGN complex for a final spindle orientation.

Epithelial cells physically interact with the surrounding extracellular matrix via integrins and with neighbouring cells via adherens junctions, providing several possible such molecular means of transmitting cues for spindle orientation^23^. As the cell rounds up in metaphase, the protrusion that mediate connections to the substrate are termed retraction fibres^13, 24–28^. Characterisation of these fibres suggests a context-dependent molecular composition at the site of tethering to the substratum, which then propagates tensile forces through the fibre^29, 30^. The molecules that provide tensile strength to the retraction fibre, and mediate the functional connection from fibre to LGN, are not yet well understood.

Subsequent to spindle orientation, membrane remodelling provides a second critical step to ensuring that daughter cells are appropriately positioned following cell division^31^. The mechanisms by which daughter cells reclaim the space vacated by their parent involve a similarly complex interplay involving adhesion complexes, tension and geometry ^32^. In epithelial monolayers, this process is complicated by the need to maintain epithelial barrier function, which necessitates continual interaction with neighbouring cells ^33^. As with spindle orientation, membrane remodelling involves a dynamic relationship between the dividing cell and its neighbours, involving physical forces and signalling adhesion proteins such as E-cadherin^33^. New filopodia are formed to facilitate cell spreading and adhesion, and these filopodia are thought to be guided by the retraction fibres that recorded previous positioning of the parent cell^15, 34^. A key focus of research has been on tricellular junctions, which are formed after cytokinesis on either side of the midbody, and can determine whether the daughter cells maintain contact after cell division, or whether these contacts are displaced by neighbouring cells^35^. The mechanisms of membrane remodelling in the absence of neighbouring cells has been less well studied.

Here, we used a simple model of single MCF10A cells dividing on plastic to explore the minimal requirements for spindle orientation, membrane remodelling and daughter cell positioning. We made the surprising finding that a complex of E-cadherin and Scribble localises at the base of retraction fibres to dictate spindle orientation, and characterized the molecular basis of that interaction. We also found that E-cadherin and Scribble subsequently relocate to the nascent junction between the two daughter cells, and orchestrate a spreading of that junction to facilitate enduring connection between the daughters. The findings described below show that Scribble traffics dynamically between key regions of the cell during cell division, coordinating the E-cadherin-mediated effects on the spindle, and extension of filopodia and the nascent daughter-daughter junction.

## Results

### E-cadherin controls cell-autonomous spindle orientation and positioning of daughter cells

We first explored the molecular composition of cell protrusions during cell division in the MCF10A human mammary epithelial cell line. During prometaphase, cells exhibited long protrusions rich in F-actin and E-cadherin, but with minimal active β1 integrin (**Fig. 1ai**). These protrusions are traditionally termed retraction fibres, and are retained as cells round up for cell division By the telophase stage, the cell contained shorter protrusions, with more active β1 integrin in the adhesions and less E-cadherin (**Fig. 1aii**, quantified in **Fig. 1aiii**). The enrichment of E-cadherin in prometaphase retraction fibres was a surprise given that E-cadherin is classically considered to mediate cell-cell interactions, rather than cell-matrix interactions^36, 37^. It also does not seem likely that E-cadherin in the retraction fibres is recruited in a ligand-dependent manner, since the substrate was not coated with E-cadherin ligands, and the E-cad in fibres was no stronger at the substrate-contacts of the fibres than along their lengths (see XV view of E-cad, Fig 1a). Further characterisation indicated that the β1 integrin-associated protein, paxillin, was also not detected in metaphase protrusions, but was clearly at the base of the protrusions by late telophase (**Supplementary Fig. 1a**). Of proteins known to mediate E-cadherin functions, β-catenin, but not β-PIX, was detected in protrusions throughout mitosis (**Supplementary Fig. 1b,c**). Together, these findings indicate that the retraction fibres of single MCF10A cells in prometaphase, rather than containing canonical integrin-based adhesions, contain E-cadherin.

**Figure 1.**
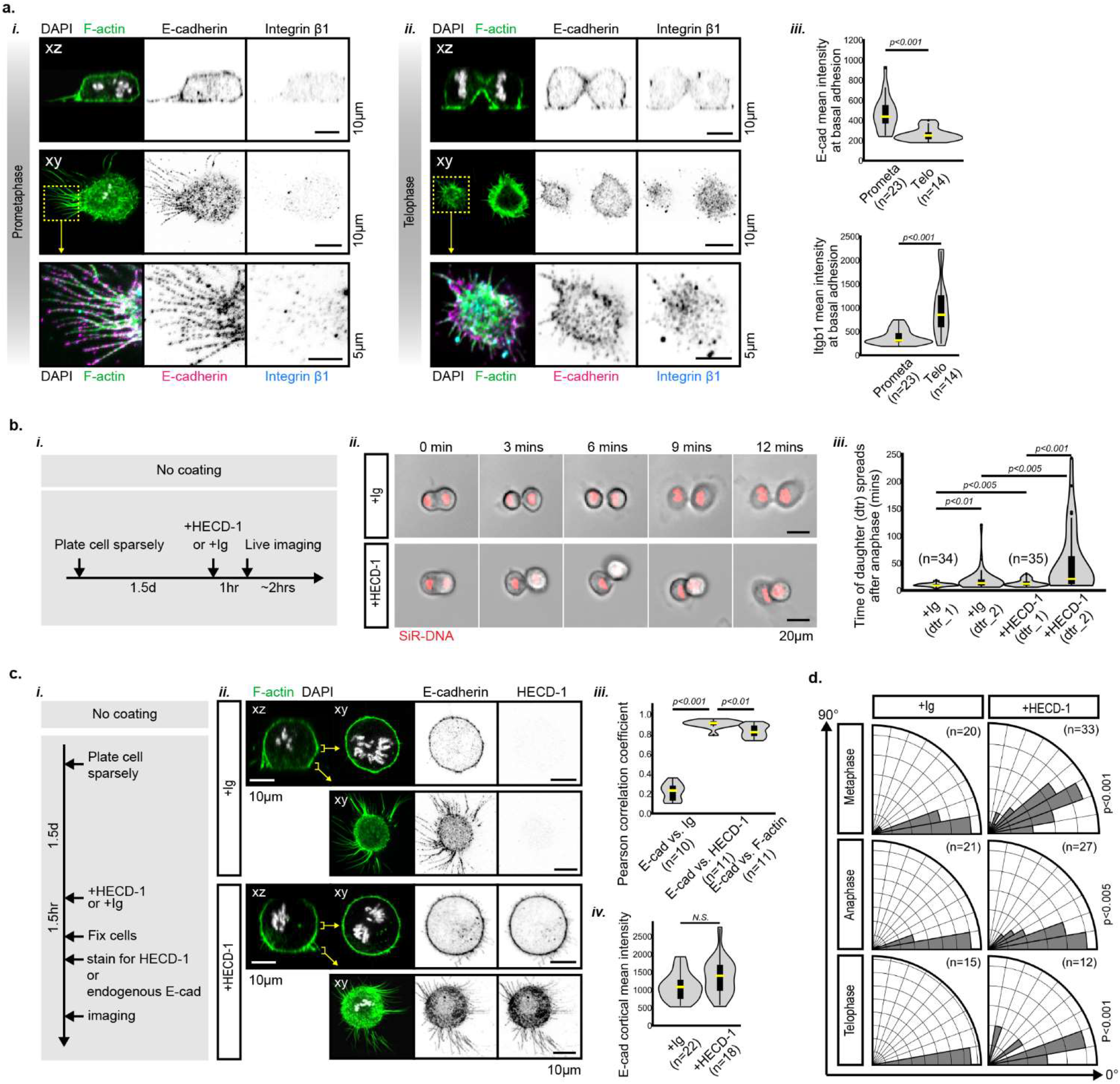
E-cadherin is required for cell-autonomous control of spindle orientation and daughter cell attachment. **a.** Sparsely plated MCF10A cells were stained with DAPI and phalloidin (pseudo-coloured white and green respectively) to label chromatin and F-actin respectively, and with antibodies to E-cadherin and active integrin β1. Representative single cells in prometaphase (i) and telophase (ii) are shown in X-Z (left) and in X-Y with two sections of the z-stack shown to represent the centre and basal regions. Enlargement of a region of interest in the basal XY slice highlights the localisation of F-actin, E-cadherin and activated integrin β1 (pseudo-coloured according to the protein labels) on the fibres linking the cell to the substrate. (ii) quantifies the average intensity of E-cadherin (upper) and activated integrin β1 (lower) in the basal adhesion region of prometaphase and telophase cells, represented by a density plot (violin plot) over laid with a box plot in which the mean/median is shown as a yellow bar, the line indicates the first and last quartile, and the box the second and third quartile, p values were calculated using a two-tailed unpaired t test **b,** MCF10A cells were labelled with SiR-DNA and treated as indicated in the schematic (i), and subsequent cell divisions were imaged by time lapse fluorescence microscopy (ii) to compare treatments with control antibody (‘Ig’) or blocking antibody to E-cadherin (‘HECD-1’). The time for each daughter cell to re-adhere to the plastic and spread was recorded as d1 (first to attach) and d2 (second daughter to attach) and shown as density plots overlaid with box plots (iii). **c**, MCF10A cells were treated as indicated in the schematic (i) to assess the localisation of HECD-1 and its impact on endogenous E-cadherin by immunofluorescent staining to discriminate between the HECD-1 antibody (anti-mouse Ig) and endogenous E-cadherin (ii). Co-localisation of endogenous E-cadherin with control Ig, HECD-1 or F-actin was assessed using Pearson correlation (iii) and the effect of HECD-1 treatment on endogenous E-cadherin intensity at the cortex was measured (iv). **d,** MCF10A cells were treated with HECD-1 and the Ig control as in **c**(i), and were stained with antibody against tubulin to measure the mitotic spindle orientation in confocal images. Spindle orientation at metaphase, anaphase and telophase were quantified and shown in polar histograms (90° indicates the spindle is perpendicular to the surface). p values were calculated using a one-tailed unpaired t test.

We explored the functional implications of E-cadherin enrichment in retraction fibres during mitosis using a blocking antibody to E-cadherin, HECD-1 that prevents E-cadherin-E-cadherin interactions^38, 39^ (**Fig. 1b**). We first explored the time taken for the daughter cells to re-adhere and spread on the substrate after cell division. In untreated cells, both daughter cells adhered to the substrate within a relatively short time. However, in the presence of HECD-1 treatment, we consistently observed that of two daughter cells produced per division, one would be significantly inhibited in its re-adherence and re-spreading, while the other was unaffected. Thus, E-cadherin is not required for spreading *per se*, but is required for optimal re-adherence and spreading of the second daughter cell after division of MCF10A cells.

To determine whether the HECD-1 antibody directly influenced the localisation of E-cadherin, we stained with a second antibody to E-cadherin after HECD-1 treatment (**Fig. 1c**). Overlays showed strong co-localisation of HECD-1 and anti-E-cadherin, indicating both that the HECD-1 antibody was specific, and that the impact of HECD-1 was not to substantially reorganise the E-cadherin localisation.

One possible explanation for these findings might be that E-cadherin is required for the mitotic spindle to be oriented parallel to the substrate during division, ensuring both daughters have access to substrate for re-adherence following mitosis. E-cadherin can regulate spindle orientation in the context of an epithelial tissue^40–43^, and it was recently shown that E-cadherin knockdown disrupts spindle orientation in an immortalized prostate epithelial cell line, RWPE-1^40^. In some contexts, spindles are randomly oriented in metaphase, and only achieve appropriate orientation in anaphase, but in other contexts, alignment occurs in metaphase^44–47^. Single MCF10A cells showed the latter behaviour, with spindles reliably aligned in metaphase, anaphase and telophase (**Fig. 1d**). However, in cells that had been treated with HECD-1, the spindle was frequently oriented away from the substrate, particularly in metaphase (**Fig. 1c**). Thus, inhibiting E-cadherin interactions leads to asymmetric adhesion of the daughter cells, associated with a mis-oriented spindle in the dividing mother cell. Interestingly, this finding contrasts with a previous finding that low focal adhesion connectivity to the substrate led to horizontal spindle orientation of HeLa cells, which was disrupted by increased focal adhesions^48^.

Together, these findings indicate that E-cadherin plays an unexpected role in transmitting information about prior adhesions to the substrate to a mitotic cell, influencing daughter cell positioning by controlling the orientation of the mitotic spindle.

### Scribble scaffolds E-cadherin to orient the mitotic spindle

How might E-cadherin influence spindle orientation in this context where intercellular junctions are not available? One clue might come from recent observations that in confluent monolayers, E-cadherin is recruited to regions of high tension, to which it directly recruits LGN^41, 42^. LGN then engages with NuMA upon breakdown of the nuclear envelope, which orchestrates pulling on astral microtubules^41, 42^. It is well established that stabilization of E-cadherin is key to its function at cell junctions, where E-cadherin stability is promoted by homophilic adhesions^49^. It is less clear what might promote stability of E-cadherin at the cell-substrate interface. One candidate is the polarity protein, Scribble. Scribble functionally interacts with E-cad, and is known to orient the mitotic spindle^40, 50–54^. Scribble and E-cad reciprocally regulate each other’s positioning and stabilization in a context-dependent manner^53, 55, 56^. Staining of MCF10A cells undergoing mitosis showed Scribble, E-cadherin and F-actin at the cortex, and in retraction fibres (**Fig. 2a**). Zooming in on the retraction fibres showed that Scribble and E-cadherin were anti-correlated, with alternating peaks of expression spaced approximately 200 nm apart along the retraction fibres. X-Z views of the dividing cells indicated that cortical Scribble, E-cadherin and F-actin were consistently enriched at the poles of the cells in metaphase (**Fig. 2b**). This position is consistent with previously observed localisation of active integrin β1 recruiting the LGN complex to control spindle orientation in single mitotic HeLa cells^13^. The position of Scribble and E-cadherin at these sites therefore provides for a scenario in which they might recruit NuMA and the LGN complex to orient the mitotic spindle.

**Figure 2.**
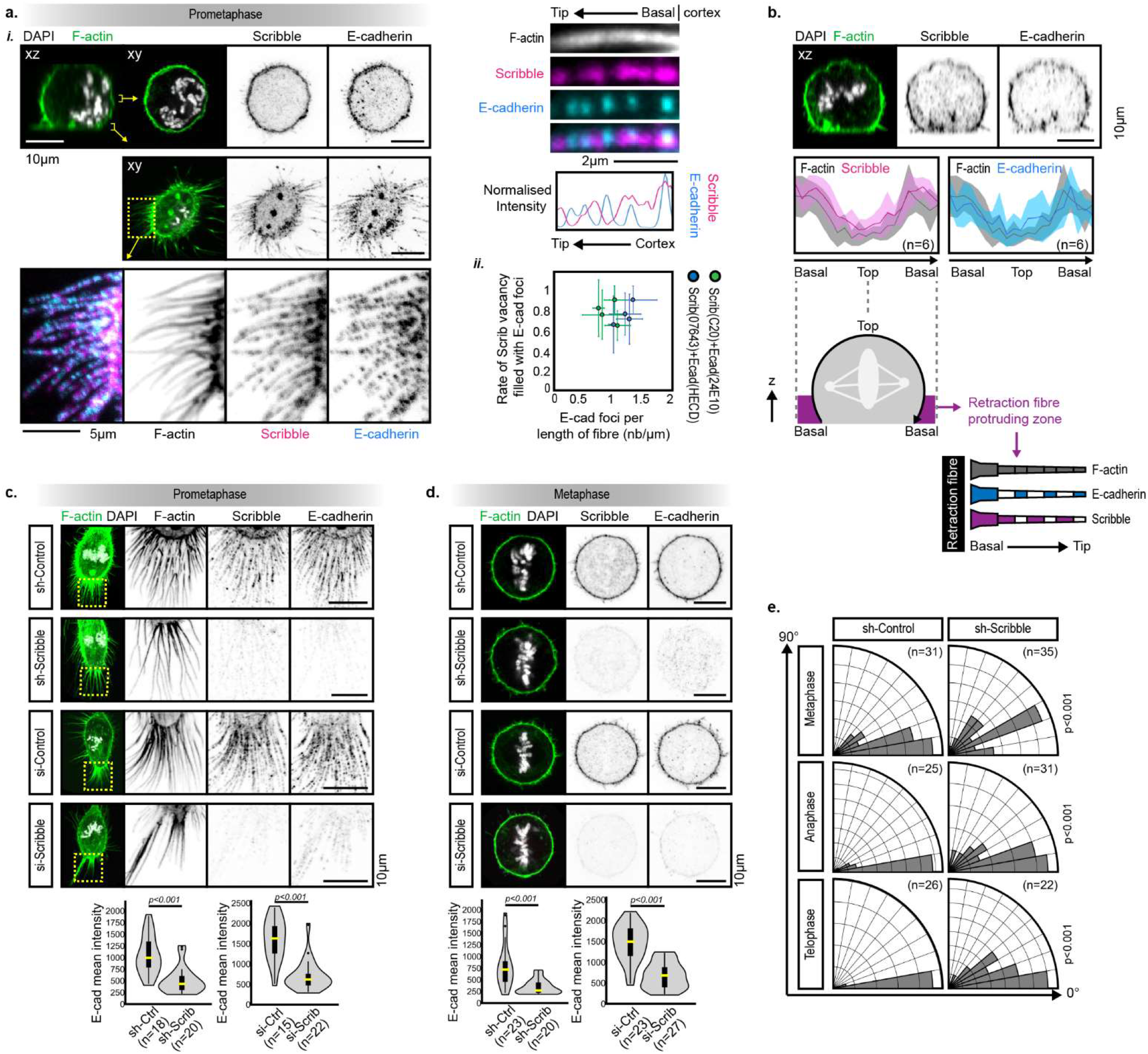
Scribble scaffolds E-cadherin at the mitotic cortex and retraction fibres to orient the mitotic spindle. **a.** Sparsely seeded MCF10A cell were co-stained for F-actin, E-cadherin, Scribble and DAPI and imaged by confocal microscopy at different stages of division. A representative image at prometaphase (i) shows Scribble and E-cadherin co-localised at the cortex (seen in the z-section from the cell centre) and in retraction fibres (seen in the basal z-section and zoomed in below). Both E-cadherin and Scribble appear punctate and alternating in the fibres, and quantification of the normalised intensity along a fibre demonstrates this alternating pattern. To determine the reproducibility of this alternating expression in the retraction fibres, the number of E-cadherin foci per µm along a fibre was plotted against the proportion of Scribble ‘vacancies’ (patches that did not contain Scribble). This alternating expression pattern in the retraction fibres is illustrated in the schematic bottom right. **b.** Scribble was depleted in MCF10A cells to assess its influence on E-cadherin localisation on the retraction fibre during prometaphase (**b)** and on the cortex during metaphase (**c**). Cells were treated with sh-Control, sh-Scribble, si-Control, and si-Scribble (rows 1-4 respectively), and the mean intensity of E-cadherin in the retraction fibres (b) or cortex (c) was shown as density plots overlaid with box plots. **d**. The effect of Scribble knockdown on spindle orientation was tested. MCF10A cells with sh-Scribble and the sh-Control were stained with antibody against tubulin to measure the mitotic spindle orientation. Spindle orientation at metaphase, anaphase and telophase were quantified and shown in polar histograms (90° indicates the spindle is perpendicular to the surface). p values were calculated using a one-tailed unpaired t test. **e**. e, Both Scribble and E-cadherin are enriched near retraction fibres of metaphase cells. Left, an xz view of a metaphase cell. The line intensity profile (right) of Scribble and E-cadherin were measured in the xz image, as the illustration (Middle) shows. (n=6 cells analysed)

To test whether Scribble impacted E-cadherin stability in either retraction fibres or the cortex, we depleted Scribble in MCF10A cells using two knockdown approaches: si-RNA and sh-RNA. Scribble was uniformly depleted by both knock-down approaches (**Supp. Fig. 2a**). Depletion of Scribble had no impact on adhesion following cell trypsinization (**Supp. Fig. 2b**), and no impact on the levels of E-cadherin at the cortex of cells in a confluent monolayer (**Supp. Fig. 2c**). However, Scribble-depleted cells showed almost complete loss of E-cadherin in retraction fibres at prometaphase (**Fig. 2c**) or at the cortex at metaphase (**Fig. 2d**). This, combined with the normal expression of E-cadherin in Scribble-depleted MCF10A cells in interphase, suggests that Scribble is required for the recruitment or maintenance of E-cadherin in the cortex and retraction fibres during cell division.

Given our finding that Scribble and E-cadherin functionally interact on retraction fibres, and the published evidence that all three members of the Scribble complex (Scribble, Discs large and Lethal Giant Larvae) mediate spindle orientation in intact tissues^40, 50, 51, 56–60^, we next investigated whether Scribble was required for spindle orientation in single cells. Indeed, in MCF10A cells lacking Scribble and undergoing cell division, the spindle was mis-oriented (**Fig. 2e**). This phenotype was similar to that observed with loss of function of E-cadherin (Fig 1c). Thus, both Scribble and E-cadherin regulate spindle positioning, and the positioning of E-cadherin in retraction fibres is dependent upon Scribble.

### Scribble coordinates NuMA positioning at the cortical poles for spindle orientation

We next explored whether Scribble and E-cadherin utilized the canonical mediators of spindle orientation^6^. Spindle orientation is dictated by dynein-mediated forces, which are generated when the microtubule motor protein, dynein, pulls on astral microtubules emanating from the spindle pole ^44, 61^. The position of the dynein-based motor complex is controlled by the position at the cortex of a complex termed the LGN complex (comprising Gαi, LGN, and NuMA in mammals). The positioning of the LGN complex is highly context specific, but often controlled by polarity proteins, so we speculated that Scribble might function by enabling the E-cadherin-based recruitment of the LGN complex to cortical poles. In wild-type single MCF10A cells, NuMA localised to spindle poles and weakly at the cortex at metaphase, but not in the retraction fibres (**Fig 3a**). This is compatible with previously described cell cycle-dependent influence of LGN on NuMA localisation^63, 64^. To determine any functional relationship between Scribble and the LGN complex, we assessed NuMA localisation during metaphase in the context of Scribble depletion. Both methods of Scribble depletion reduced the cortical recruitment of NuMA at metaphase (**Fig. 3b**). The specificity of this effect is demonstrated by our observation that Scribble depletion did not impact the localisation of Myosin IIb (**Supp. Fig. 2d**).

**Figure 3.**
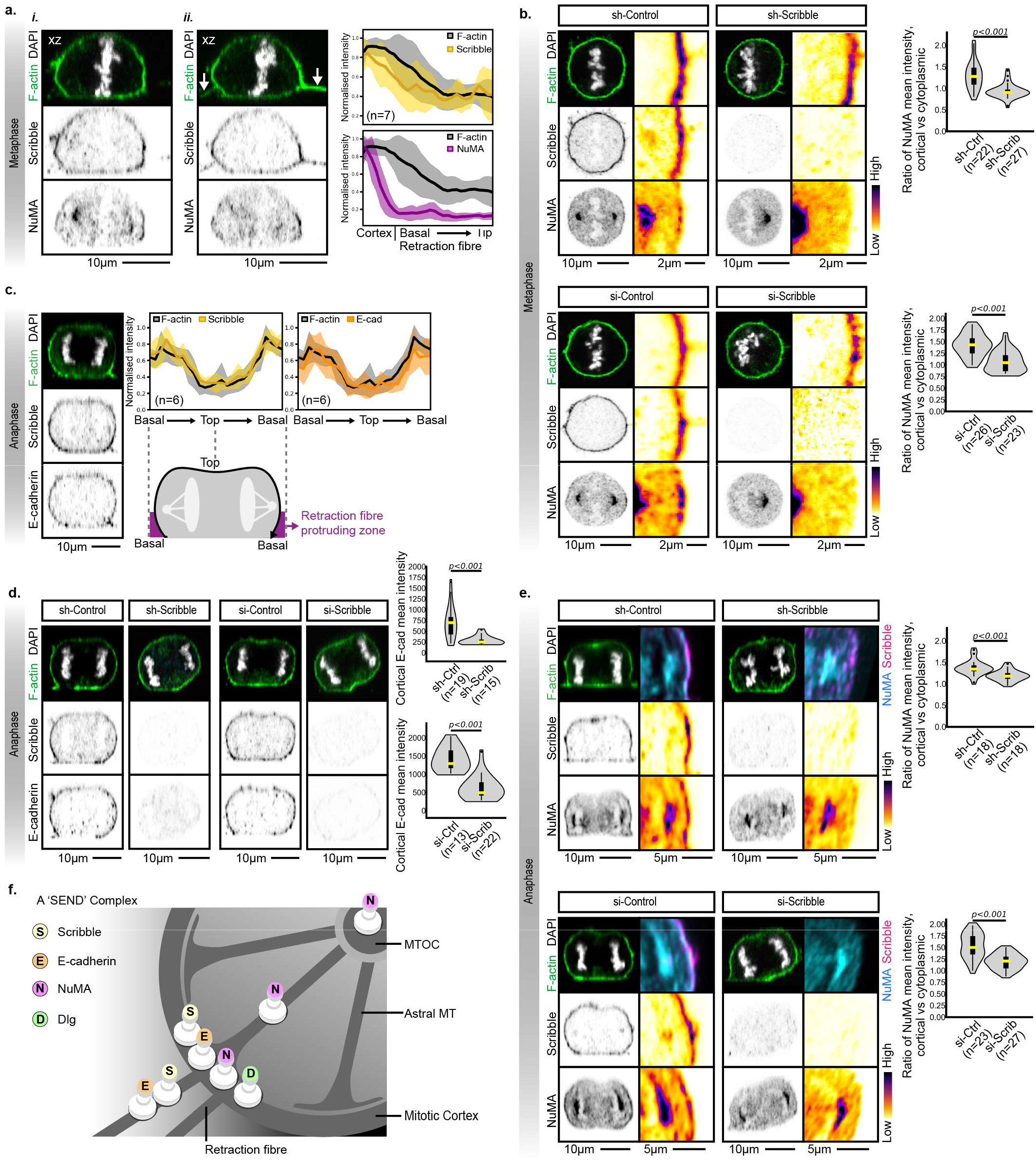
Scribble is a scaffold for NuMA at the cortex during metaphase and anaphase. **a**, To assess whether Scribble and NuMA co-localized during cell division, sparsely seeded MCF10A cells were stained with DAPI (DNA), phalloidin (F-actin) and antibody against Scribble and Paxillin. **(i)**, In a representative metaphase cell, two XZ planes are shown, one of an XZ plane at the cell centre illustrating that Scribble and NuMA co-localised at the retraction fibre attachment region, and one plane that contained retraction fibres, showing that Scribble, but not NuMA was expressed in the retraction fibres. **b**. A schematic to suggest a ‘SEND’ complex (<u>S</u>cribble, <u>E</u>-cadherin, <u>N</u>uMA, <u>D</u>lg) at the mitotic cortex bridges the signals that passes through retraction fibre to the astral microtubule for spindle positioning. **c**. MCF10A cells were depleted of Scribble using shRNA and si-RNA as shown, to assess the impact on NuMA localisation during metaphase. The ratio of cortical to cytoplasmic NuMA was shown as density plots overlaid with box plots (p value, two-tailed unpaired t test) and showed reduced localisation at the cortex with both means of depletion. **d**. To assess if the SEND complex might contribute in later mitotic phases, cells stained as in (a) were captured in anaphase. Both Scribble and E-cadherin are enriched at the retraction fibre attachment region of the anaphase cells. Left, an XZ view of a representative anaphase cell; Right, line intensity profile of Scribble and E-cadherin in 6 cells. **e**. Scribble was depleted in MCF10A cells by both sh-RNA and si-RNA, and E-cadherin localisation to the anaphase cortex was assessed as shown in density plots overlaid with box plots (p value, two-tailed unpaired t test). **f.** MCF10A cells were depleted of Scribble using shRNA and si-RNA as shown, to assess the impact on NuMA localisation during anaphase. The ratio of cortical to cytoplasmic NuMA was shown as density plots overlaid with box plots (p value, two-tailed unpaired t test) and showed reduced localisation at the cortex with both means of depletion.

Scribble played similar roles in anaphase as we had observed in metaphase cells. As previously reported^64^, NuMA levels are dramatically increased at anaphase (**Supp. Fig 3**), and, similarly to metaphase, at anaphase E-cadherin and Scribble co-localised to the cortical pole (**Fig. 3c**). Again, depletion of Scribble using either siRNA or shRNA prevented cortical localisation of E-cadherin at anaphase (**Fig. 3d**). Similarly, depletion of Scribble reduced cortical NuMA at anaphase (**Fig. 3e**). These results suggest that cues from retraction fibres are transmitted through Scribble and E-cadherin to recruit NuMA to the cortical pole at the base of the fibre for spindle orientation (**Fig. 3f**).

To determine whether the action of Scribble was dependent upon E-cadherin we tested HeLa cells, which lack E-cadherin expression (**Supp. Fig. 4**)^65^. Similar to MCF10A cells, NuMA recruitment to the cortex at anaphase was also dependent upon Scribble expression in HeLa cells. The mitotic spindle of HeLa cells on plastic was consistently oriented along the substrate, with a mild loss of control in the absence of Scribble. These data show that Scribble does not require E-cadherin to mediate NuMA recruitment to the cortex.

Together, these findings indicate that the presence of E-cadherin and Scribble in retraction fibres at metaphase leads to recruitment of NuMA to the cortical pole.. These findings are reminiscent of previous findings that Dlg, a key functional partner of Scribble^56^ , can co-localise with Scribble at the spindle pole^58^, where Dlg recruits the LGN complex to orient the mitotic spindle in several animal models and tissue types^66–68^. We therefore propose that the E-cadherin and Scribble in retraction fibres establish a ‘SEND’ complex of Scribble, E-cadherin, NuMA and perhaps Dlg (Fig. 3f). Based upon extensive literature on the role of NuMA in LGN-mediated regulation of mitotic spindle orientation, and our findings of defective spindle orientation in metaphase, anaphase and telophase in the absence of Scribble, we propose that the SEND complex, by coordinating pulling forces from the two cortical poles, coordinates stable alignment of the spindle parallel to the substrate.

### Scribble relocates to the daughter-daughter contact to control nascent junction formation

The aberrant spindle orientation observed in dividing cells with compromised Scribble or E-cadherin provides one possible explanation for the failure of the second daughter to adhere and spread (Fig 1b), but we also identified a second possible explanation. In staining the dividing cells we observed that in late telophase, Scribble relocated from the cortical pole to the region between the two nascent daughters. Scribble was not evident at the basal adhesion region between the two daughters where both F-actin and paxillin were enriched, but was strongly expressed at the daughter-daughter contact site above the intracellular bridge^69, 70^ (**Fig 4a**). We compared the localisation of Scribble, F-Actin and Myosin IIb along regions of furrow ingression and daughter-daughter contact sites at progressive telophase stages (**Supp. Fig. 5**). This suggested that, although all three proteins were broadly recruited to this region throughout cytokinesis, Scribble was co-enriched with F-Actin at the daughter-daughter contact rather than with Myosin IIB. However, activity of both was required, as treatment with Cytochalasin D and Blebbistatin abrogated recruitment of Scribble to the daughter-daughter interface (**Supp. Fig. 6**).

**Figure 4:**
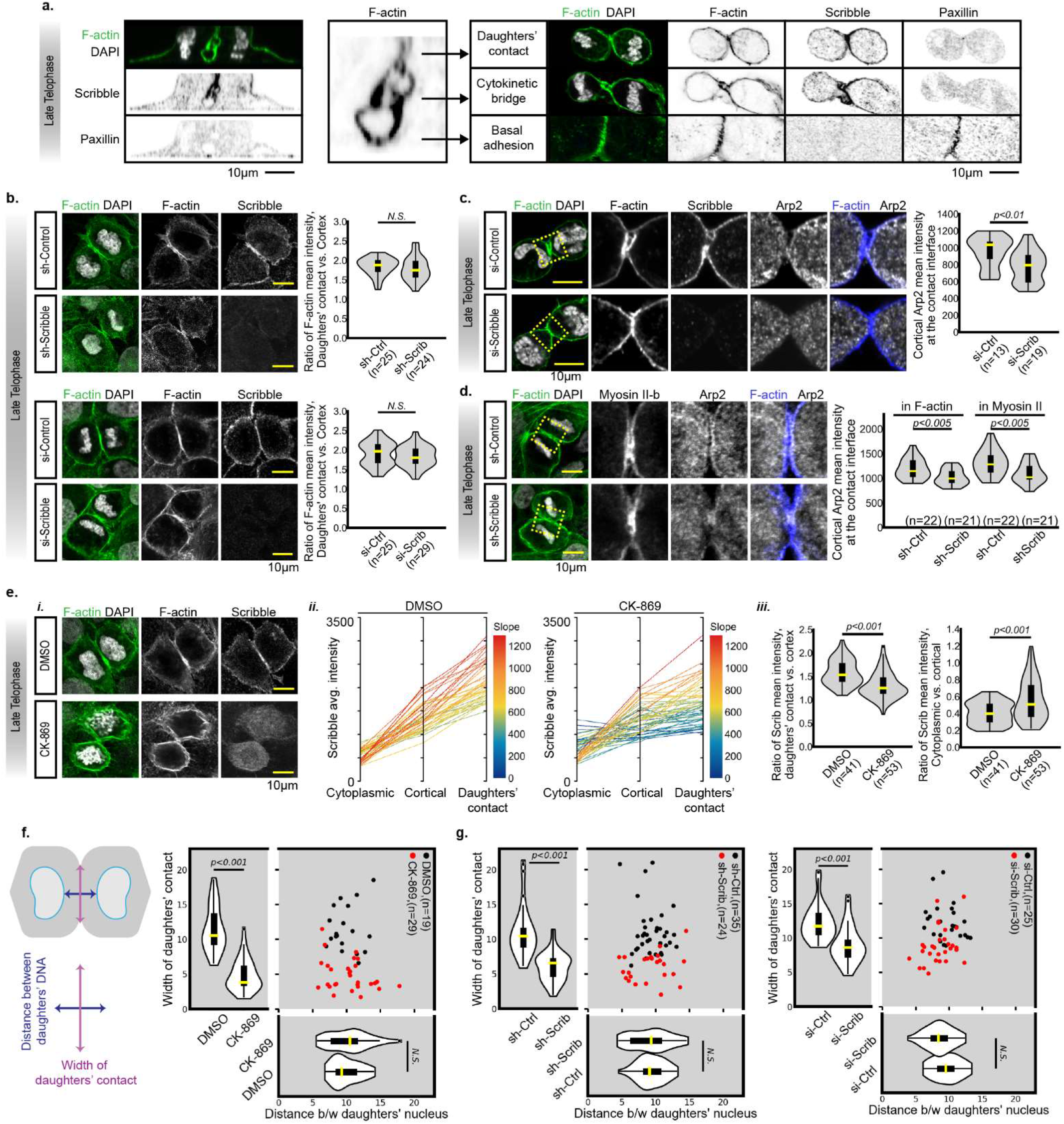
Scribble interacts with Arp2/3 and promotes elongation of the nascent daughter-daughter contact. **a,** To assess the localisation of Scribble at late telophase, sparsely plated MCF10A cells were stained with DAPI (DNA), phalloidin (F-actin) and antibody against Scribble and Paxillin. The first column (left) shows an xz view sectioned through a representative cell centre. In the F-actin channel (middle), the regions of each subcellular structure (ie. ‘daughters’ contact’, ‘cytokinetic bridge’, and ‘basal adhesion’) were defined. For each region, the images stained with DAPI (DNA), phalloidin (F-actin), Scribble and Paxillin were displayed respectively (right). **b,** Scribble was depleted in MCF10A cells by both sh-RNA and si-RNA, and F-actin enrichment at the daughters’ contact was assessed as shown in density plots overlaid with box plots (p value, two-tailed unpaired t test). **c,** Scribble was depleted in sparsely plated MCF10A cells by si-RNA, and Arp2 localised at the cortical region of daughters’ contact was assessed as shown in density plots overlaid with box plots (p value, two-tailed unpaired t test). **d,** Scribble was depleted in MCF10A cells by sh-RNA, and Arp2 localised at the cortical region of daughters’ contact was assessed as shown in density plots overlaid with box plots (p value, two-tailed unpaired t test). **e,** To investigate the role of Arp2/3, sparsely plated MCF10A cells were treated with Arp2/3 inhibitor CK-869 (50μ) for 15 minutes, and stained with DAPI (DNA), phalloidin (F-actin) and antibody against Scribble. A representative image is shown in (i), and the mean intensity of Scribble at subcellular localisation, including cytoplasm, cortex, and daughters’ contact was quantified in (ii) and the ratio in (iii). For the absolute Scribble intensity analysis in (ii), the lines connect data points from individual cells, with the colour spectrum highlighting the slope of each line, indexed by red to high slope and blue to low slope. In (ii), the ratio of Scribble mean intensity at cytoplasm or daughters-daughter contact versus cortex were further shown in density plots overlaid with box plots (p value, two-tailed unpaired t test). **f,** Sparsely plated MCF10A cells treated with Arp2/3 inhibitor CK-869 as in (e) were assessed for the width of daughter-daughter contact and the distance between daughter cell chromatin as per the schematic (Left), and were shown in the scatter plot and density plots overlaid with box plots (p value, two-tailed unpaired t test). **g,** Scribble-depleted MCF10A cells as in (b) were assessed similarly for the width of daughter-daughter contact and the distance between daughter nuclei were analysed as the method illustrated in (f), and were shown in the scatter plot and density plots overlaid with box plots (p value, two-tailed unpaired t test).

We assessed whether the presence of Scribble at the late telophase daughter-daughter contact site might reflect a role in establishing stable connections between the daughters. We found no difference in the intensity of F-actin at the contact site when cells were depleted of Scribble (**Fig. 4b**). However, Arp2/3, a key mediator of actin branching that enables daughter-daughter adhesions in the context of Drosophila dorsal thorax epithelial cells^71, 72^ was reduced at the daughter-daughter contact in Scribble-depleted cells (**Fig. 4c**). This reduction in Arp2/3 was seen in both the F-actin-rich and Myosin IIb-rich regions of the daughter-daughter interface (**Fig. 4d**). These data suggest that Scribble is not required for recruitment of F-actin to the daughter-daughter interface, but might facilitate the actin branching mediated by Arp2/3, perhaps to stabilize or extend this interface. A similar role for pushing of actin protrusions for E-cadherin adhesion was recently identified in interphase MDCK cells^71, 73^. Treatment with the Arp2/3-specific inhibitor, CK-869, reduced both the overall cortical recruitment of Scribble and the proportion of cortical Scribble that was concentrated at the daughter-daughter interface (**Fig. 4e**). These data indicate that Apr2/3 and Scribble are recruited to the daughter-daughter interface at telophase in a co-dependent manner.

### The nascent daughter-daughter interface impacts upon subsequent positioning of the daughters

We hypothesized that this recruitment to the interface might influence subsequent daughter-daughter connections. To assess this, we quantified the extent (‘width’) of the interface, and the shortest distance between the DNA (a surrogate for cell positioning) of each daughter, after Arp2/3 inhibition or Scribble depletion. Inhibition of Arp2/3 and depletion of Scribble dramatically reduced the extent of daughter-daughter contact, but had little or no impact on the cell positioning during telophase (**Fig. 4f, g**). These data indicate that actin and myosin-dependent positioning of Arp2/3 and Scribble at the interface between two daughters at telophase enables a stabilization or lengthening of the nascent daughter-daughter contact region.

To assess whether this nascent daughter-daughter contact region dictates subsequent spreading of each daughter, we first assessed whether, as with E-cadherin inhibition (Fig. 1b), impeding this contact might influence the re-adherence to the substrate of one or both daughters after cell division. Apr2/3 inhibition did not significantly influence the time taken for either daughter to re-adhere (**Fig. 5ai, ii**), but over 30 minutes, the distance between daughter cells diverged, with the distance significantly increased with Arp2/3 inhibition (**Fig. 5aiii**). This finding supports the notion that Arp2/3-mediated expansion of the daughter-daughter adhesion zone might retain the daughters in proximity to each other. Depletion of Scribble led to a slight increase in the time taken for daughter 1 to re-adhere to the substrate, and a dramatic increase for daughter 2 (**Fig. 5bi, ii**). Indeed, some of the second daughters failed to re-adhere before they were lost to the tracking, perhaps because the spindle mis-orientation reduced association with the substrate. The distance between daughter cells could only be quantified for those that adhered within 30 min, and showed no difference (**Fig. 5biii**). However, the lack of effect of Scribble on daughter cell distance should be treated with caution given the issues with delayed re-adherence in Scribble-depleted cells. Overall, these data indicate that the timing and or position of daughter cell re-adhesion after cytokinesis is influenced by both Scribble and Arp2/3 activity.

**Figure 5:**
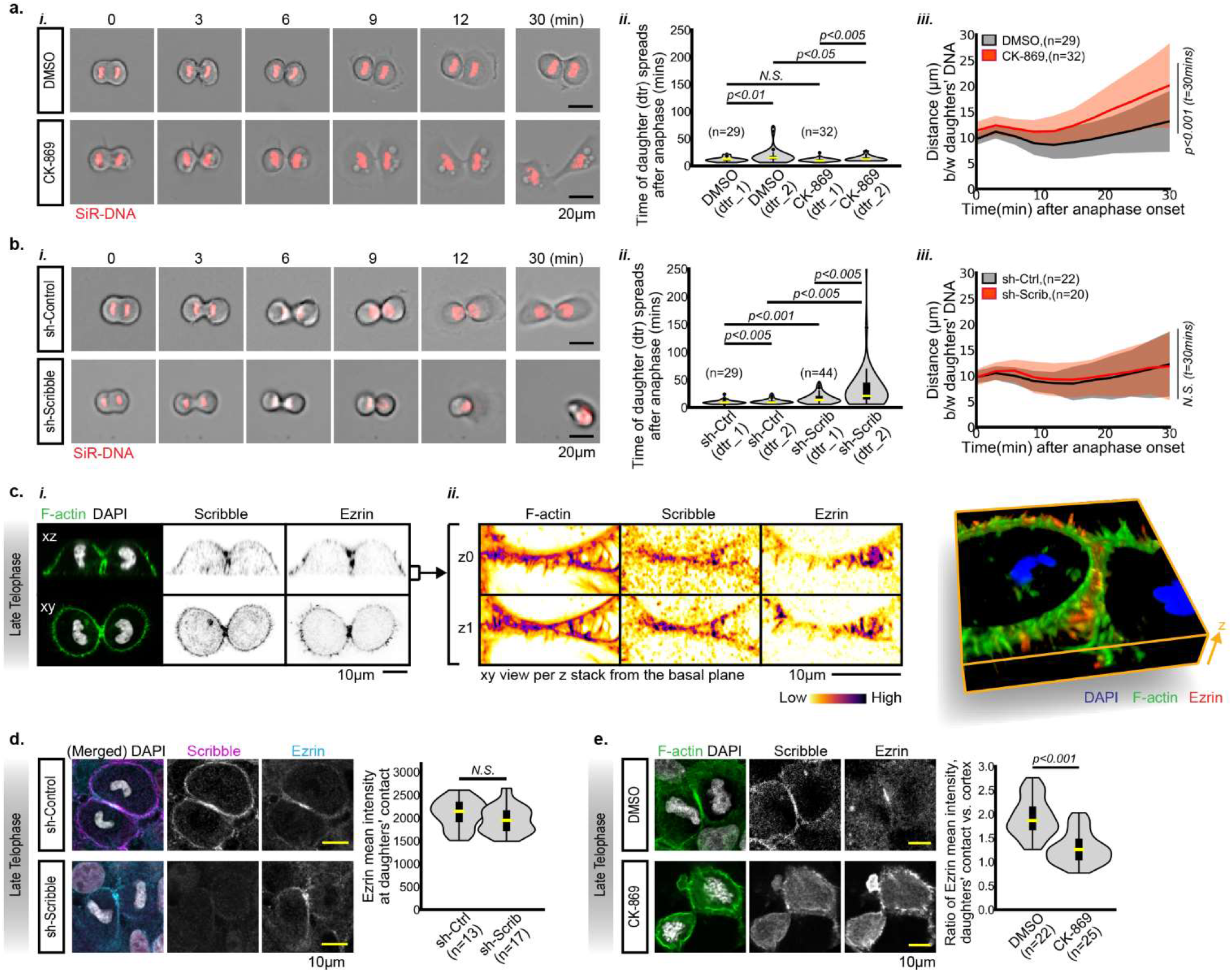
Arp2/3 cooperates with Scribble and Ezrin to govern daughter cell geometry during cell division. **a,** To assess the role of Arp2/3 in daughter cell positioning, MCF10A cells were labelled with SiR-DNA (shown in red), treated with DMSO control or inhibitor CK-869 (0.1 μM), and imaged by time lapse microscopy following division (i). The time for each daughter cell to re-adhere to the plastic and spread was recorded as d1 (first daughter to attach) and d2 (second daughter to attach) and shown as density plots overlaid with box plots (ii). The distance between daughters’ nucleus was measured from the anaphase onset (time = 0 minute) for 30 minutes, and was shown in the line plot, with shadow represented one standard deviation (iii). **b,** To assess the role of Scribble in daughter cell positioning, analysis as per (a) was performed on MCF10A cells with sh-RNA knockdown control (‘sh-Control’) or Scribble knockdown (‘sh-Scribble’). **c,** The protein localisation of Scribble and Ezrin at late telophase was assessed in a sparsely plated MCF10A cell stained with DAPI (DNA), phalloidin (F-actin) and antibody against Scribble and Ezrin. A representative image (i) is shown as sequential Z stacks (3 per image) in (ii), and in a 3D reconstructed view (iii). **d,** The role of Scribble in Ezrin recruitment to the daughter-daughter interface was assessed by comparing MCF10A cells with sh-Scribble and sh-Control (representative image on the left, quantification on the right, p value, two-tailed unpaired t test). **e,** The role of Arp2/e in Ezrin recruitment to the MCF10A daughter-daughter interface was assessed using the inhibitor CK-μM) 15 minutes, and the mean intensity of Ezrin enriched at the daughter-daughter contact was assessed as shown in density plots overlaid with box plots (p value, two-tailed unpaired t test).

We further explored the nature of the nascent daughter-daughter junction, and identified intercalated filopodia that were rich in the membrane-cytoskeletal linker, Ezrin, Scribble, and F-actin (**Fig. 5c**). Scribble-depleted cells showed no obvious defects in Ezrin at the daughter-daughter interface (**Fig. 5d**). However, Ezrin at the interface was dependent upon Arp2/3 activity (**Fig. 5e**), similar to the dependence of Scribble on Arp2/3 activity. Together, these experiments suggest a model in which Scribble, Ezrin and Arp2/3 are recruited to interdigitations at the nascent junction, where they enable expansion of the nascent daughter-daughter contact and positioning of one daughter relative to the other.

### The composition of filopodia evolves throughout cell division, and Scribble influences nascent post-mitotic filopodia

A role of retraction fibres is thought to be the marking of cell position, such that nascent filopodia can reclaim the tracks of past filopodia^15, 34^. To assess any molecular and functional relationship between retraction fibres and post-mitotic filopodia in our system, we first further characterised the filopodia at different stages of cell division. The extensive protrusions observed in prometaphase gradually disappeared, with few observed in anaphase (**Supp Fig 7**). We first explored Myosin X, an unconventional myosin that couples actin-dependent forces from retraction fibres to centrosomes^74^, and is recruited to nascent focal adhesions in some contexts^75, 76^. Zooming in on individual filopodia indicated that Myosin X was present in discrete puncta along the length of the single MCF10A retraction fibres of prometaphase cells, and these puncta overlapped with Scribble puncta (**Fig. 6a**). In contrast, Myosin X was predominantly present in at the tips of filopodia during telophase and late telophase, compatible with these representing nascent adhesions. Scribble localisation did not overlap with that of Myosin X in these nascent filopodia. In contrast to Mysoin X, the membrane-cytoskeletal adaptor, Ezrin was excluded from the cortex, and highly expressed at the base of prometaphase retraction fibres, and the mid-section of nascent filopodia (**Fig. 6b**). The change in composition between the pre-anaphase retraction fibres and the post-anaphase nascent filopodia is described in **Fig 6c**.

**Figure 6:**
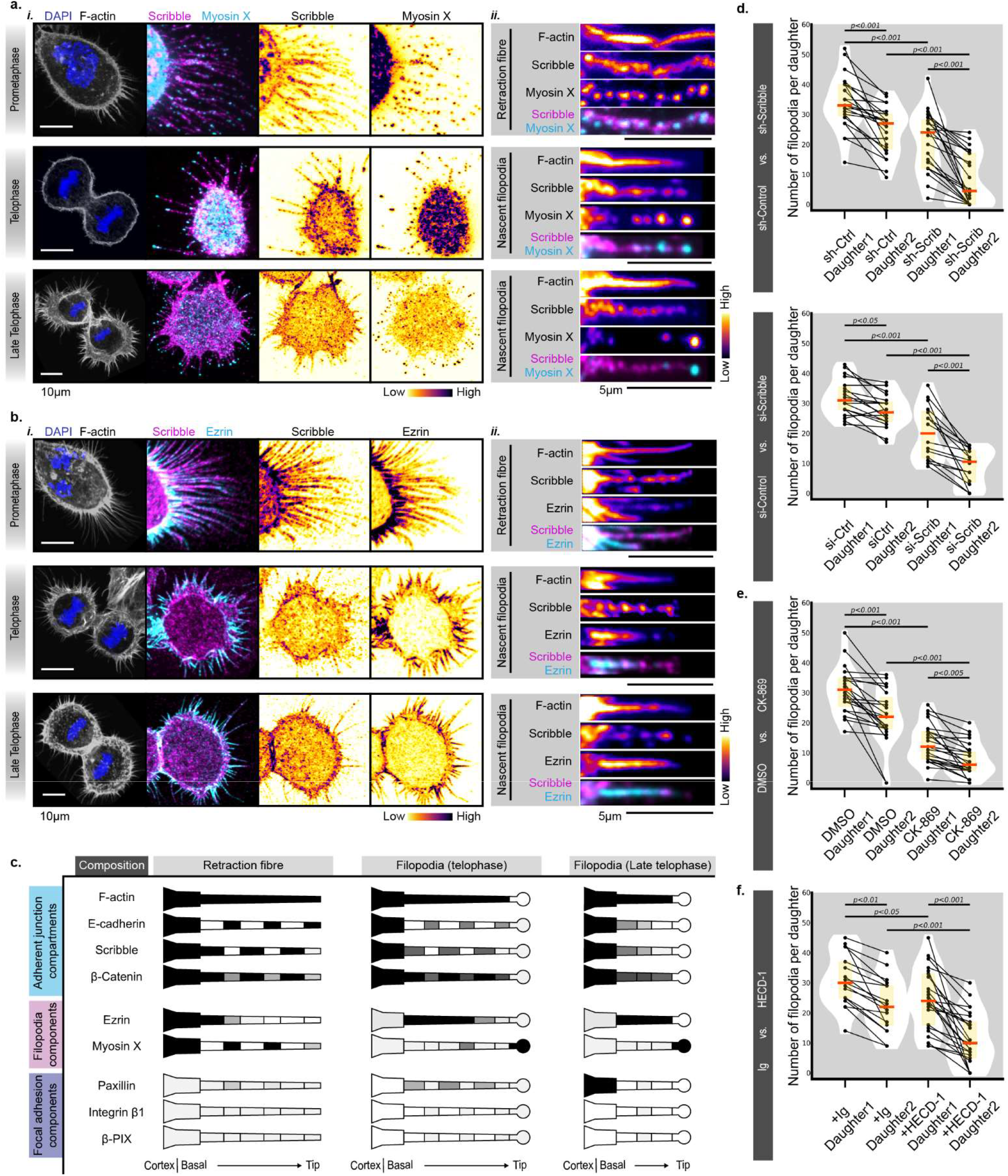
Scribble is required for nascent filopodia patterning. **a,** The localisation of Scribble and Myosin X was assessed at prometaphase, telophase and late telophase (Left), with DAPI (DNA), phalloidin (F-actin) co-labelling. The fire coloured images (Right) were the regions of retraction fibre or nascent filopodia cropped from the left images. **b,** Protein localisation of Scribble and Ezrin at prometaphase, telophase and late telophase (Left), with DAPI (DNA), phalloidin (F-actin) co-labelling. The fire coloured images (Right) were the regions of retraction fibre or nascent filopodia cropped from the left images. **c,** A schematic to summarise the protein localisation of F-actin, E-cadherin, Scribble, β-catenin, Ezrin, Myosin X, Paxillin, Integrin β and β-PIX at retraction fibre and filopodia. The colour of black represents high protein intensity, the white represents the protein intensity was nearly undetectable, and the grey represents the protein intensity is in between the black (high) and the white (none). **d, e, f,** The sparsely plated MCF10A cells, with the treatments (see Methods) of Scribble knockdown (‘sh-Scrib’ and ‘si-Scrib’), knockdown control (‘sh-Ctrl’ and ‘si-Ctrl’), Arp2/3 inhibition (‘CK-869’, 50μM for 15 minutes), (‘DMSO’), E-cadherin blockade (‘HECD-1’), Ig control (‘Ig’), were labelled with F-actin, and the filopodia protruded from the basal adhesion of each daughter cell were counted and shown in the line plot, overlaid with the density plot and the box plot (p value, two-tailed unpaired t test).

If the components of the retraction fibres are instrumental in production of nascent filopodia, one would expect a functional effect beyond that predicted by the spindle misorientation defects described above. We therefore assessed the number of nascent filopodia in post-mitotic MCF10A cells depleted of Scribble, focussing specifically on the cell divisions where both daughters had re-attached. It has long been appreciated that post-mitotic adhesion is dependent upon filopodia^77^, so we assessed the number of filopodia separately for the daughter that adhered first (’daughter 1’) and the daughter that adhered second (’daughter 2’. We examined the cells immediately upon flattening onto the substrate (**Fig. 6d**). Indeed, both Scribble-depleted daughters exhibited a significant reduction in the number of filopodia at late telophase. A similar reduction in filopodia from both daughters was observed after treatment with the Arp2/3 inhibitor (**Fig 6e**), confirming the importance of Arp2/in filopodia remodelling. Interestingly, inhibition of E-cadherin led to reduced nascent filopodia in the second daughter (compatible with reduced spreading due to mis-oriented spindle) but had a less convincing effect on the number of nascent filopodia in the first daughter. Together, these data suggest the possibility that Scribble, and perhaps E-cadherin, regulate the transition from retraction fibre to nascent filopodia, providing opportunities to reposition the daughter cells according to the position of the cell prior to cytokinesis.

### Scribble coordinates an E-cadherin-based nascent junction to dictate daughter cell positioning

Given the collaboration between Scribble and E-cadherin in coordinating spindle orientation at the metaphase stage, we explored a functional interaction in the formation of the nascent junction at telophase. Indeed, E-cadherin was co-enriched with Scribble at the nascent daughter-daughter junction (**Fig 7a**), and as we previously observed for Scribble, recruitment of E-cadherin and Myosin IIb required Arp2/3 activity (**Fig. 7b**). Interestingly, depletion of Scribble resulted in a dramatic loss of E-cadherin, and this was directly correlated with the degree of Scribble depletion ((**Fig. 7c**). As we had previously observed, the effect of Scribble depletion on E-cadherin expression was not observed in a confluent monolayer of MCF10A cells in interphase (Supp Fig 2), so we now assessed E-cadherin in the nascent junction of cells dividing within a confluent monolayer, using Myosin IIb as a marker of the mature telophase junction^36^. Interestingly, like the single cells, E-cadherin was recruited to the nascent daughter-daughter junction within a confluent monolayer, and this recruitment was dependent upon Scribble (**Fig 7d**). Given that Dlg generally co-operates with Scribble^56^ and has previously been found at nascent daughter-daughter contacts^78^, we propose the acronym SEAD as a functional complex that mediates expansion of the nascent daughter-daughter contact. (**Fig. 7e**). This expanded daughter-daughter junction then restricts the movement of the two daughters away from each other.

**Figure 7:**
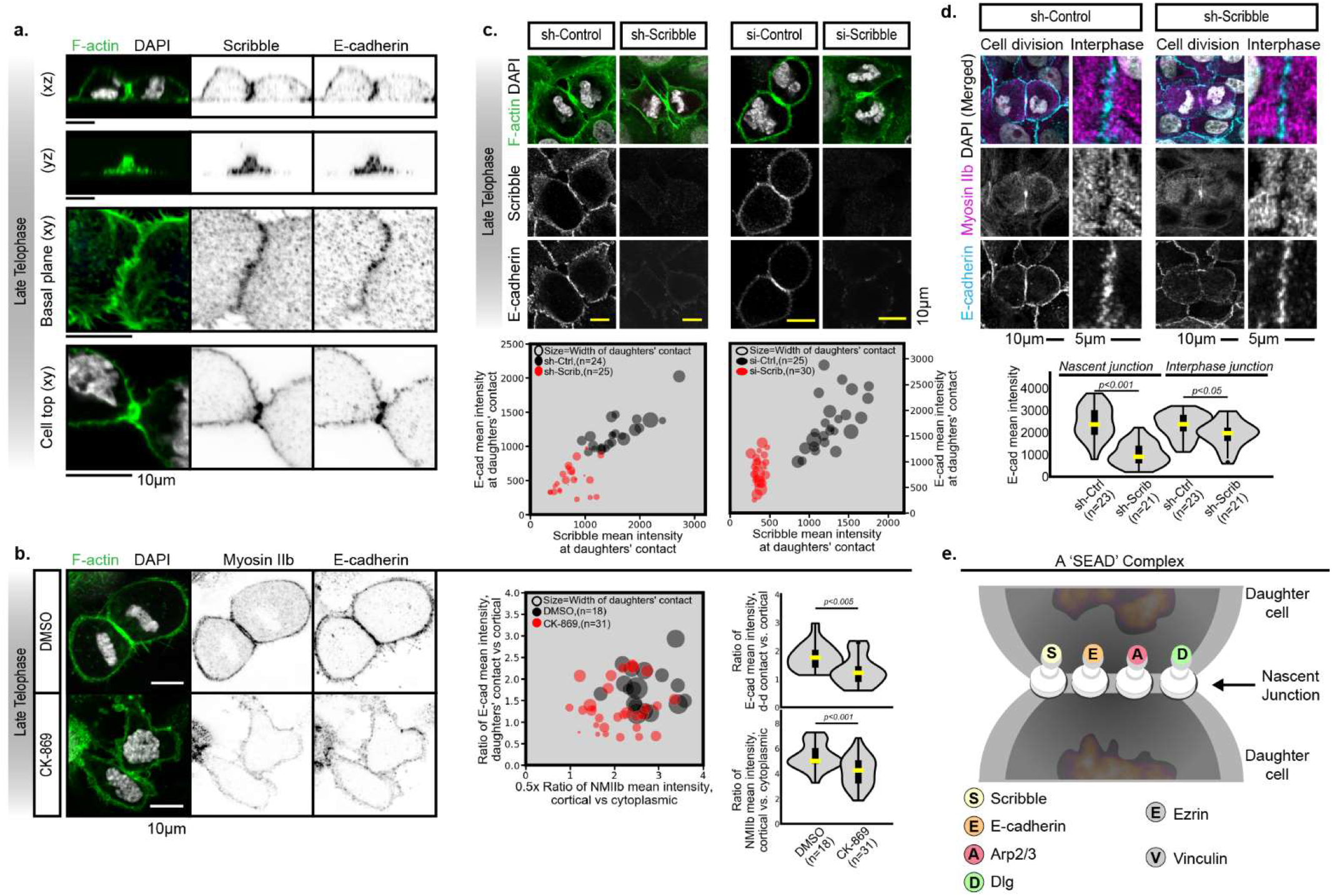
Scribble and E-cadherin regulate the expansion of a nascent junction between daughters. **a,** Protein localisation of Scribble and E-cadherin at late telophase. Representative image shows a sparsely plated MCF10A cell stained with DAPI (DNA), phalloidin (F-actin) and antibody against Scribble and E-cadherin. The (xz) layout shows the view that sectioned through the cell centre. The (xz) layout shows the view of the interface of daughter-daughter contact. The two (xy) layouts show the view at the basal adhesion and the top of the cell respectively. **b,** Inhibitor CK-869 (50 μM) was treated on MCF10A cells for 15 minutes. (Left) Representative image shows a sparsely plated MCF10A cell stained with DAPI (DNA), phalloidin (F-actin) and antibody against Myosin IIb and E-cadherin. Ratio of mean intensity of Myosin IIb at cortex versus cytoplasm, and ratio of mean intensity of E-cadherin enriched at daughters’ contact versus cortex were co-plotted in a scattering plot (Middle), with size of per dot determined by the width of daughters’ contact. Based on the same measurement, the density plots overlaid with box plots (p value, two-tailed unpaired t test) were shown (Right). **c,** Scribble was depleted in MCF10A cells by both sh-RNA and si-RNA, and mean intensity of Scribble and E-cadherin were assessed as shown in the scattering plots below, with size of per dot determined by the width of daughters’ contact. **d,** Scribble was depleted in MCF10A cells by sh-RNA, and mean intensity of E-cadherin at the nascent junction between daughters or at the interphase junction was assessed as shown in density plots overlaid with box plots (p value, two-tailed unpaired t test). **e,** A schematic to suggest a ‘SEAD’ complex (<u>S</u>cribble, <u>E</u>-cadherin, <u>A</u>rp2/3, perhaps <u>D</u>lg) regulates the nascent junction between daughters.

## Discussion

By exploring MCF10A cells without neighbours, we have identified a novel cell-autonomous mechanism of controlling spindle orientation and daughter cell positioning. Our experiments reveal a ligand-independent role for E-cadherin in transmitting signals from retraction fibres to NuMA of the LGN complex. In contrast to the situation during interphase, the stability and position of E-cadherin is profoundly dependent on Scribble during cell division. We also show that remodelling during cell division leads to further functions for E-cadherin and Scribble, in both the formation of nascent filopodia, and the expansion of a nascent daughter-daughter junction, which controls the positioning of both daughters at and beyond telophase (**Figure 8**).

**Figure 8:**
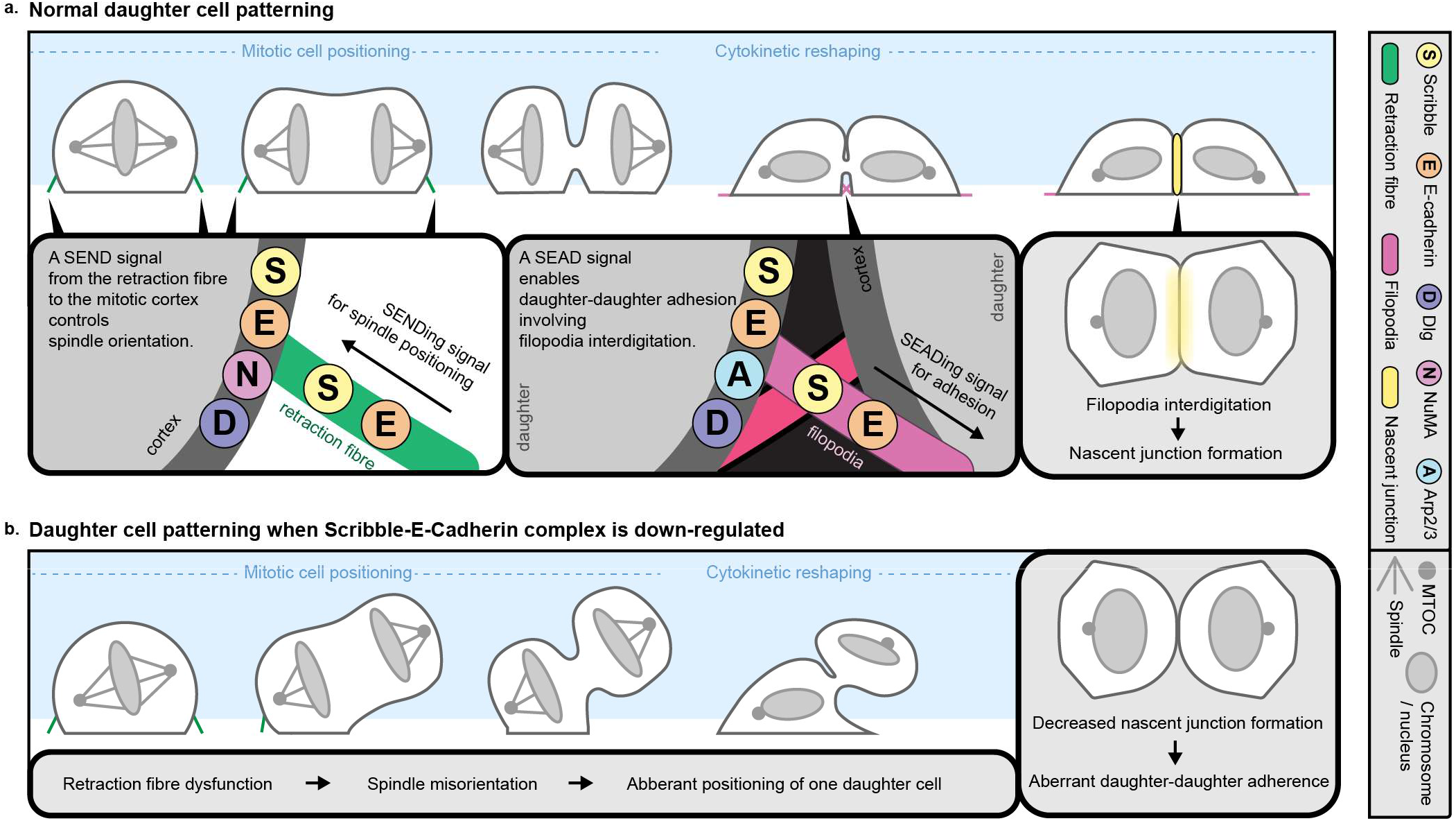
A model of daughter cell patterning controlled by a Scribble-E-cadherin functional complex. **a.** Cell shape during cell division is patterned by stepwise programs, involving spindle orientation leading to the aberrant position of one daughter cells, followed by formation of a nascent daughter-daughter adhesion. The two programs are controlled by signalling platform involving Scribble and E-cadherin. In the case of spindle positioning, NuMA is also involved (a ‘SEND’ complex) and for the nascent daughter-daughter adhesion, Arp2/3 is also involved (a ‘SEAD’ complex). **b**. A normal progression of daughter cell patterning is controlled by SENDing the signal for spindle orientation through retraction fibres, and then by SEADing the drivers for daughter-daughter adhesion. **c**. Daughter cell patterning in absence of SENDing or SEADing signals are abnormal, such as biased mitotic cell positioning, aberrant daughter-surface adhesion and reduction of daughter-daughter re-adherence.

### SENDing signals from retraction fibres to orient the mitotic spindle

E-cadherin is well known to orient the mitotic spindle by transmitting signals from cell neighbours via adherens junctions^41, 42^. Similarly, integrins can orient the mitotic spindle by transmitting signals from the cell matrix via retraction fibres^15^. In single MCF10A cells without artificial matrix, we were surprised to find E-cadherin, rather than integrin-β1, communicating from the retraction fibres. This finding adds to the growing appreciation that multiple receptors can input signals from the extracellular matrix to control mitotic spindle orientation, including β1-integrin in cells plated on fibronectin, and β5 integrin in U2OS cells that make their own extracellular matrix^80^.

Provision of artificial ligand on the adhesion surface has previously been shown to orchestrate spindle orientation^13, 43, 81^, but in our system, there is no apparent ligand available, suggesting a ligand-independent role for E-cadherin. Indeed, coating of the substrate with recombinant E-cadherin as a homophilic ligand on the basal surface of cells has previously been reported to lead to spindles oriented away from the substrate^43^ A precedent for ligand-independent orienting of the spindle is that mediated by integrin β1. In support of a similar ligand-independent role for E-cadherin, cis-clustering (also known as lateral clustering) of E-cadherin can transmit signals under conformational constraints associated with membrane immobilization^82–84^. Cells use cadherin-based tension sensing to monitor and respond to the stiffness of their environment^37, 85, 86^, and E-cadherin dilution (which could be driven by stretching of the retraction fibres) can mediate force trasnduction^32^. Together with our findings, these observations suggest that the recruitment of E-cadherin and Scribble allows for transmission of tension-related information from the retraction fibres to orchestrate the spindle.

The role of Scribble in spindle orientation is well established^40, 50–53^. However, the complexity of the tissues in which this role has been investigated mean that it is still not clear what extrinsic or intrinsic cues are transmitted by Scribble^50, 51^. Our data suggests a possible effect of Scribble beyond mediating E-cadherin effects, since we observed a role for Scribble in spindle orientation in E-cadherin-deficient cells such as HeLa cells. One explanation for this comes from recent findings of an alternative means of aligning the spindle to the history of pre-mitotic cell shape involves a dynamic, oscillatory feedback between chromosomes and LGN along the short cell axis^14^. It will be interesting to assess whether Scribble plays a role in this process. Here, we show that Scribble can directly link positional information from retraction fibres to spindle orientation control. In metaphase, this role is clearly via transmission of signals from E-cadherin.

### SEADing information to guide to guide positioning of the daughter cells

Despite the recognised importance of positioning of the daughter cells following mitosis for tissue organisation during development, metastasis and other pathologies, the means by which adhesive contacts are remodelled following cell division are still unclear^87^. This understanding is particularly complicated by the need for cells in a monolayer to disengage adherens junctions with neighbouring cells alongside formation of a new adhesive interface^88^. As with the spindle orientation, exploring single cells has allowed us to identify novel mechanisms by which the daughter-daughter contact is established, although our findings suggest these mechanisms are also conserved in an intact monolayer. We find that a SEAD complex of Scribble, E-cadherin and Arp2/3 and perhaps Dlg is recruited to the intracellular bridge that remains in late telophase, where they mediate an expansion of the adhesive interface between the two daughter cells.

Interestingly, tricellular junctions can contain Scribble, Dlg and E-cadherin^89, 90^, and Dlg in tricellular junctions of the Drosophila dorsal thorax epithelium mediates disentanglement of daughter and neighbour cells and formation of the nascent daughter-daughter interface^91^. Our findings are also reminiscent of a previously identified role of E-cadherin, Arp2/3 and myosin in the formation of a nascent junction in *Drosophila* dorsal thorax epithelial cells, although in that context neighbouring cells were required to achieve a long interface^72^. Arp2/3 is also required for expansion of the nascent daughter-daughter contact of *Drosophila* sensory organ precursors in a Rac-dependent manner^92^. Given the functional association of Rac1 and Scribble in membrane protrusions^93, 94^, it is tempting to speculate that Scribble might also be required in this context. Combining our findings and the literature therefore suggests a model in which the SEAD complex plays a general role in post-mitotic membrane remodelling..

Together, these studies identify cell-autonomous roles for Scribble and E-cadherin at several stages in the remodelling of spindle and cell membranes during cell division of a single MCF10A cell. These roles were previously obscured, presumably because of the importance of Scribble and E-cadherin in the junctions of epithelial cells in monolayers. Scribble connects E-cadherin (and perhaps other cues) to spindle orientation machinery, cell protrusions and the nascent daughter-daughter contact. This dynamic repositioning of Scribble to coordinate cell morphology during times when the cell is undergoing rapid shape change has precedence in other biological situations^52, 56, 95, 96^. Thus, we propose a dynamic role for Scribble throughout cell division in orchestrating the placement of daughter cells.

## Supporting information

Methods: Antibody table

## Supplemental Material

Supplemental Material includes Online Methods, Supplemental Figures 1-7, and Table 1 which describe antibodies used in the studies.

## Acknowledgements

We thank Michal Milgrom-Hoffman, Krystle Lim, and Rebecca Stephens for technical assistance and helpful discussions, and Mirren Charnley and John Lock for helpful comments on the manuscript. This work was funded by the Australian Research Council (FT0990405 to SMR), the National Health and Medical Research Council (APP1099140 to SMR), and a Swinburne University Postgraduate Research Award to ASC. This work was done on Wurundjeri land of the Kulin nation, and we pay our respects to the Elders past and present.

## Author Contributions

ASC contributed to conceptualization, data curation, programming, formal analysis, investigation, methodology, validation, visualization, writing, reviewing and editing; YC contributed the original experiments that inspired the project; TK contributed critical advice and reagents; POH contributed with provision of reagents, discussions of the research, review and editing of the manuscript; SMR contributed to conceptualization, formal analysis, funding acquisition, methodology, project administration, supervision, visualization, writing, review and editing.

## Data Availability

All data supporting the findings of this study are available from the corresponding author on reasonable request.

## Code Availability

No custom software or algorithms were used in this study.

## Supplementary Figure legends

**Supplementary Figure 1.**
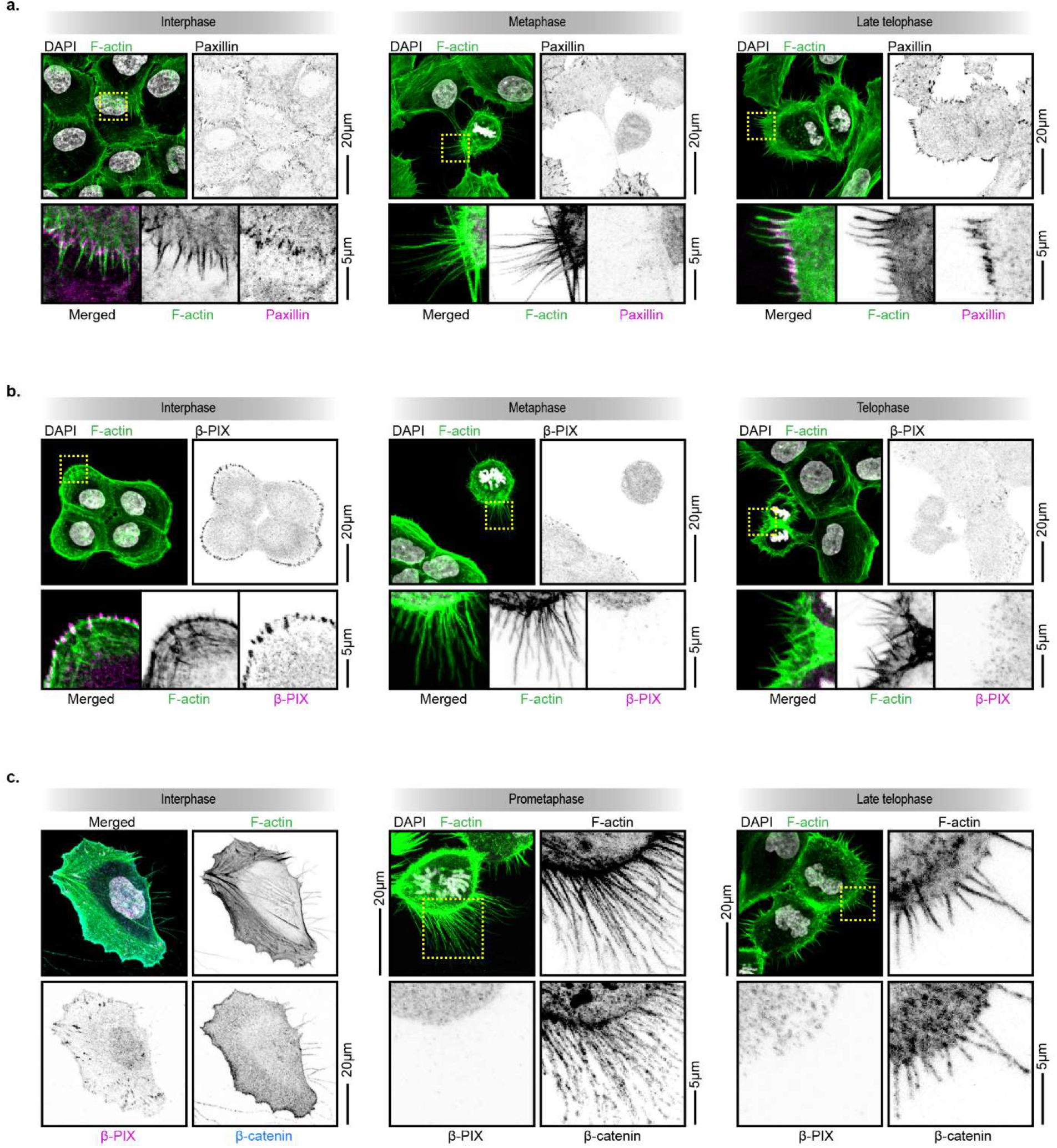
Characterisation of adhesions at different cell cycle stages of MCF10A cells. **a.** Cells at interphase, metaphase and late telophase, labelled with Paxillin, phalloidin (F-actin) and DAPI (DNA), are shown, with zoomed-in images below showing Paxillin expression in interphase filopodia, mitotic retraction fibre and cytokinetic filopodia. **b.** Cells at interphase, metaphase and telophase, labelled with β-PIX, phalloidin (F-actin) and DAPI (DNA), are shown, with zoomed-in images below showing β-PIX expression in interphase filopodia, mitotic retraction fibre and cytokinetic filopodia. **c.** Cells at interphase, metaphase and late telophase, labelled with β-catenin, β-PIX phalloidin (F-actin) and DAPI (DNA), are shown, with zoomed-in images below showing β-catenin and β-PIX expression in interphase filopodia, mitotic retraction fibre and cytokinetic filopodia.

**Supplementary Figure 2.**
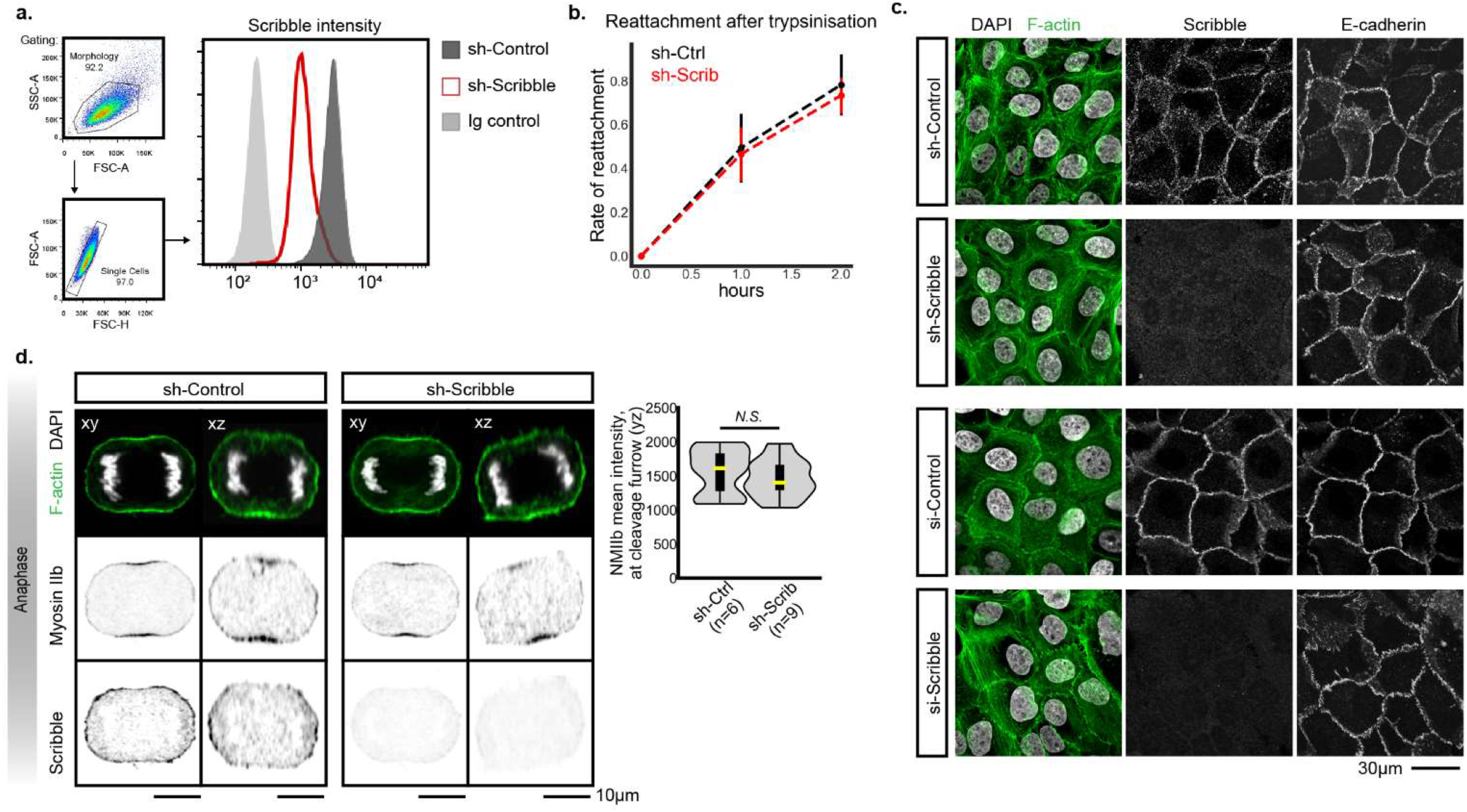
Characterisation of Scribble depletion in MCF10A cells. **a.** Flow cytometry analysis to examine Scribble depletion using sh-RNA. The cumulative histogram shows the intensity profile of intracellular staining of Scribble of the sh-Scribble cell (red), the sh-control cell (dark grey), and a control of Ig staining (light grey). The gating strategy is described in the pseudo-coloured dot plots (LHS). **b.** To examine whether Scribble depletion results in a general loss of cell adhesion, the kinetics of reattachment after trypsinisation of sh-Scribble and sh-Control cells was assessed by counting the proportion of cells that had attached at 1 and 2 hours after trypsinization. **c.** Images of interphase monolayers of MCF10A cells (sh-Scribble, sh-Control, si-Scribble and si-Control), labelled with Scribble, E-cadherin, phalloidin (F-actin) and DAPI (DNA), show that depletion of Scribble did not result in loss of interphase E-cadherin. **d.** MCF10A cells depleted of Scribble showed no difference in the recruitment of Myosin IIb to the cleavage furrow (quantified on the RHS as density plots overlaid with box plots (p value, two-tailed unpaired t test).

**Supplementary Figure 3.**
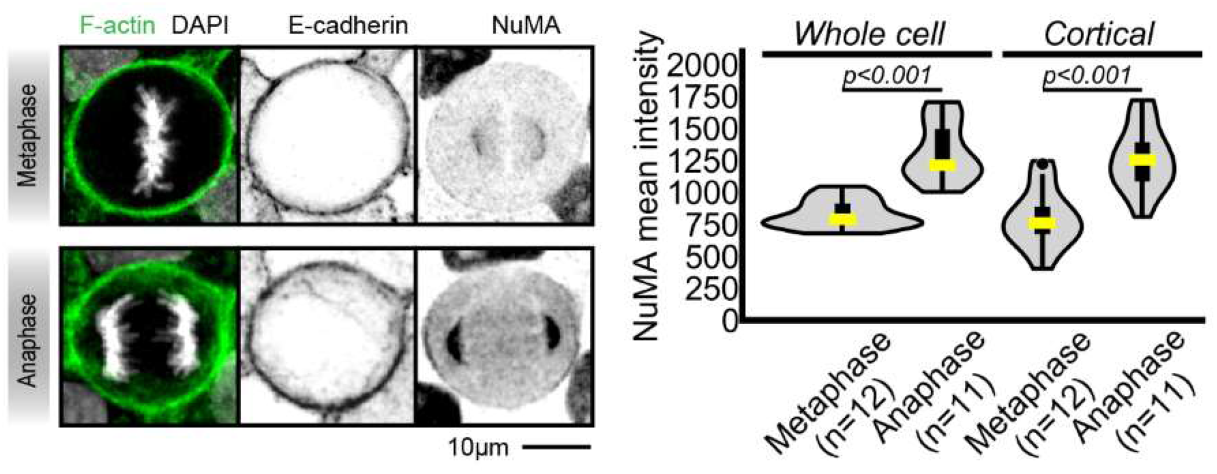
Examination of NuMA expression in MCF10A cells during metaphase and anaphase. Representative images of a metaphase and an anaphase cell, labelled with NuMA, E-cadherin, phalloidin (F-actin) and DAPI (DNA) are shown. Mean intensity of NuMA from total cell or from the cortical region was plotted in a density plot overlaid with box plot (p value, two-tailed unpaired t test).

**Supplementary Figure 4.**
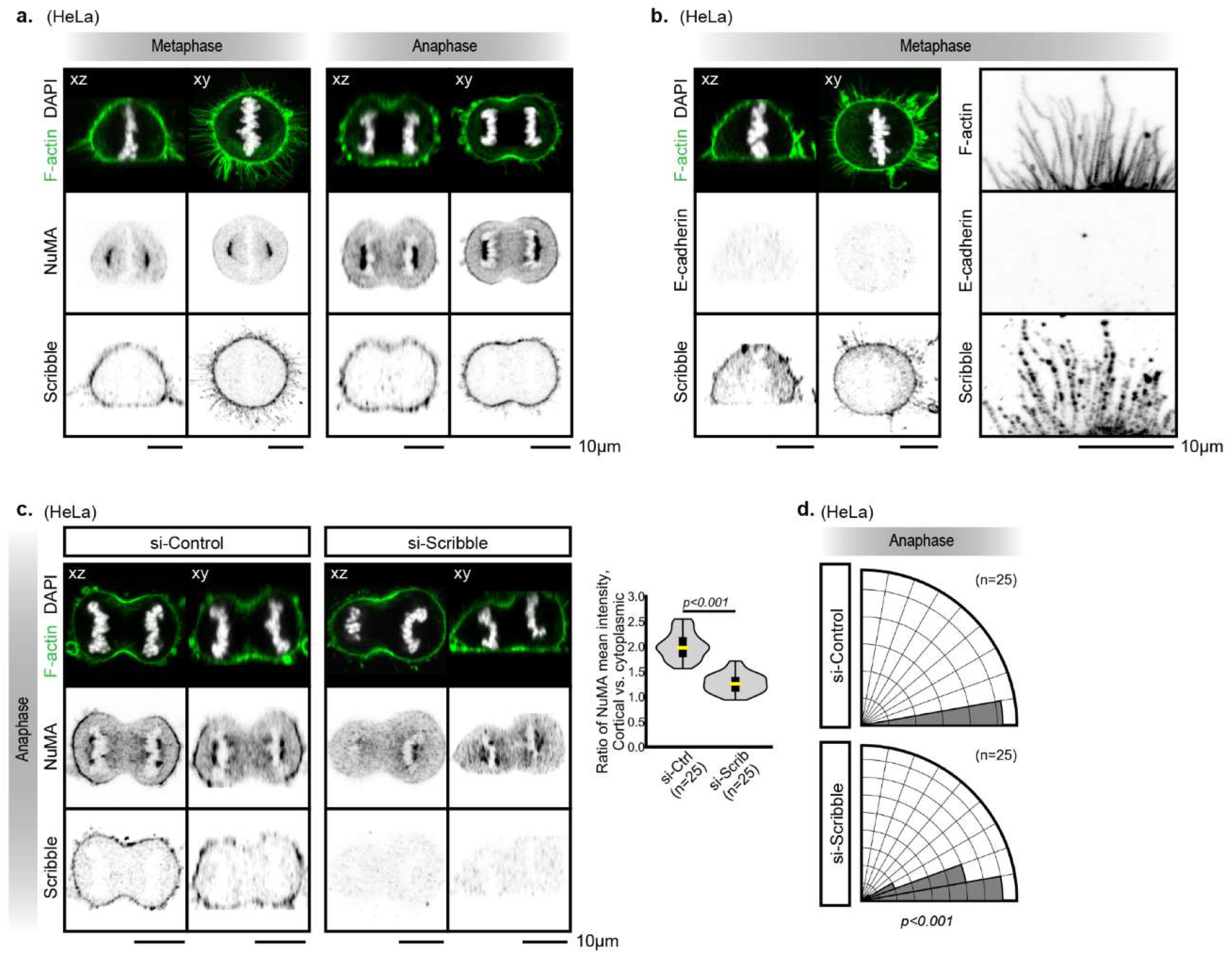
Evaluating the role of Scribble in spindle orientation in HeLa cells. **a.** Images show HeLa cells at metaphase and anaphase, labelled with NuMA, Scribble, phalloidin (F-actin) and DAPI (DNA). **b.** Images show HeLa cells at metaphase, labelled with Scribble, E-cadherin, phalloidin (F-actin) and DAPI (DNA). The zoomed-in images show the region of retraction fibre. **c.** Scribble was depleted in HeLa cells using si-RNA, and the subsequent influence on NuMA cortical localisation was assessed and plotted in density plot overlaid with box plot (p value, two-tailed unpaired t test). **d.** Scribble was depleted in HeLa cells using si-RNA, and the subsequent influence on spindle orientation was assessed and plotted in polar histograms. (p value, one-tailed unpaired t test).

**Supplementary Figure 5.**
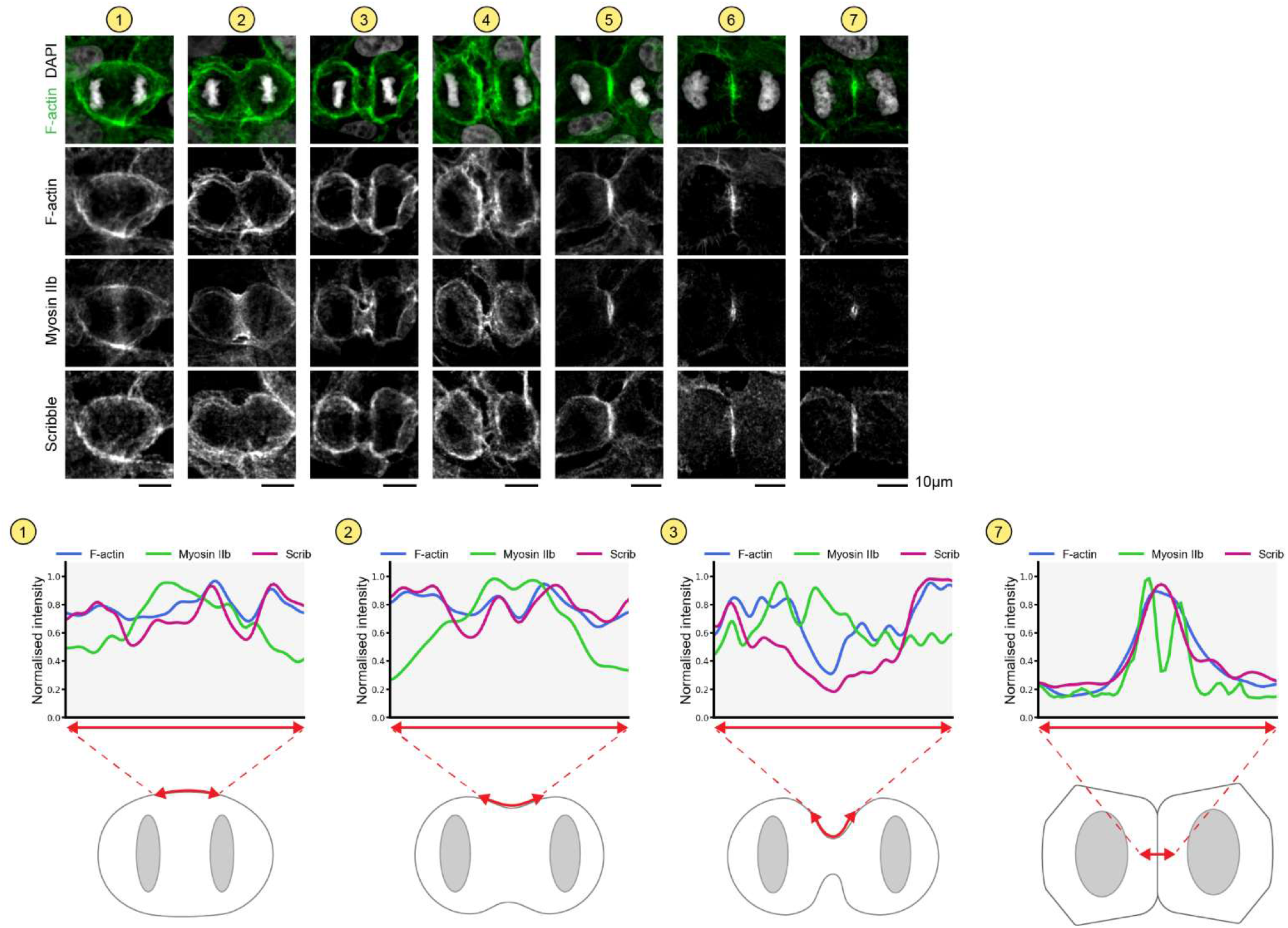
Characterisation of cortical distribution of Scribble and actomyosin during cell division. Confocal images (1∼7) show MCF10A cells at different stages from anaphase onset. Protein distribution of Scribble, Myosin IIb, and F-actin (phalloidin) at stage (1, 2, 3, 7) were assessed at the region of interests as the red line marked on the cartoon cells below.

**Supplementary Figure 6.**
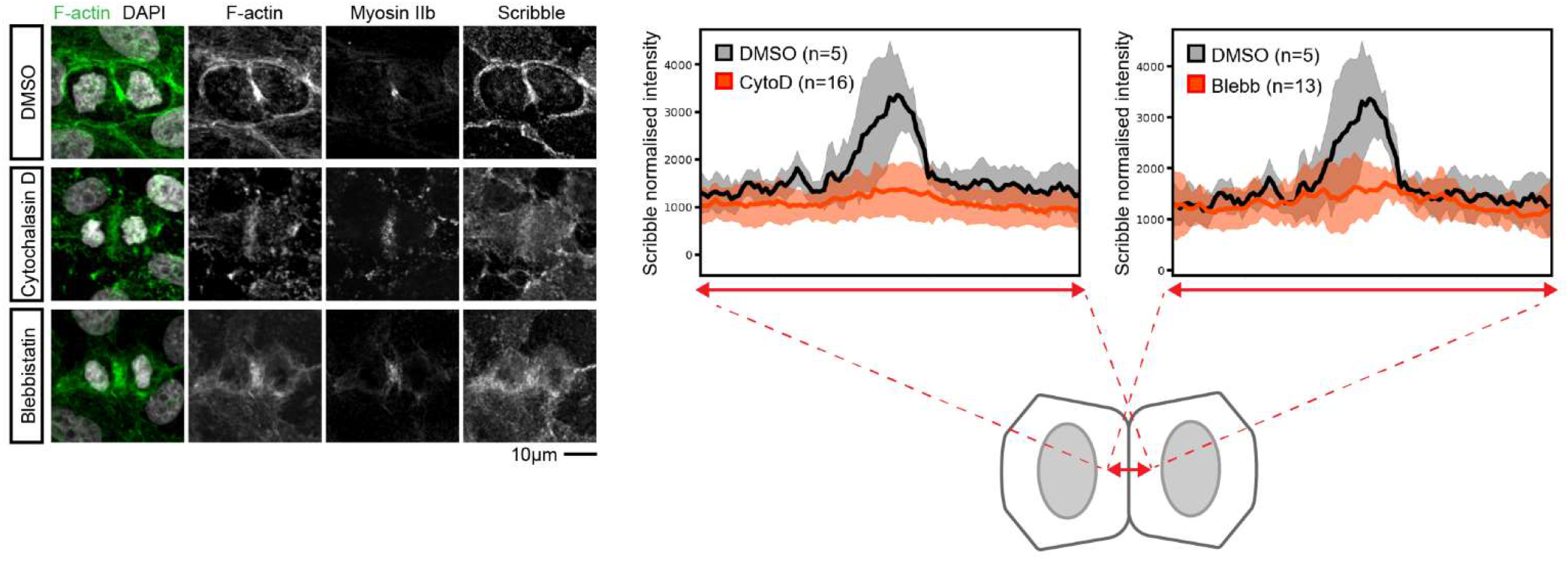
Treatment with Cytochalasin D and Blebbistatin prevent recruitment of Scribble to the nascent daughter-daughter junction. MCF10A cells were treated with drugs (DMSO, Cytochalasin D and Blebbistatin, respectively) for 15 minutes, fixed and labelled with Myosin IIb, Scribble, F-actin (phalloidin) and DAPI (DNA). The intensity profile of Scribble, measured accordingly to the red line as the schematic illustrated, were shown in the line plot in a manner of mean±SEM. Cell numbers: Cytochalasin D (n=16), Blebbistatin (n=13), DMSO (n=5).

**Supplementary Figure 7.**
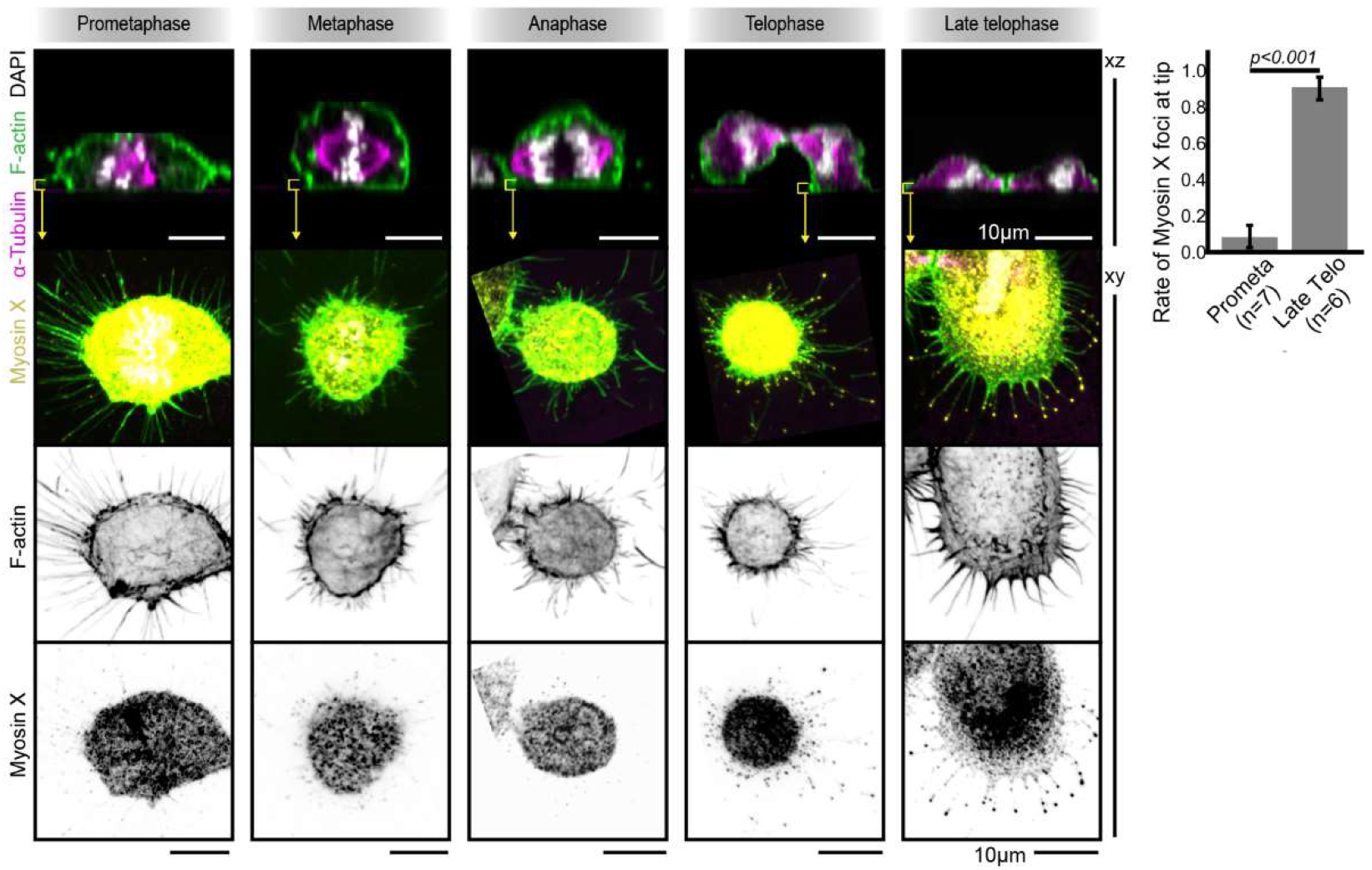
Characterisation of Myosin X distribution on cell-surface adhesion and mitotic protrusions. Confocal images show MCF10A cells, labelled with Myosin X, α-tubulin, phalloidin (F-actin) and DAPI (DNA) at the stage of prometaphase, metaphase, anaphase, telophase and late telophase. The Z-sections incorporating cell-surface adhesion were shown for each stage.

**Supplementary Table 1.** A list of antibodies and dilutions used in this study.

## References

1. Biggins, J.S., Royer, C., Watanabe, T. & Srinivas, S. Towards understanding the roles of position and geometry on cell fate decisions during preimplantation development. Semin Cell Dev Biol 47-48, 74–79 (2015).

2. Dekoninck, S. et al. Defining the Design Principles of Skin Epidermis Postnatal Growth. Cell 181, 604–620.e622 (2020).

3. van Leen, E.V., di Pietro, F. & Bellaiche, Y. Oriented cell divisions in epithelia: from force generation to force anisotropy by tension, shape and vertices. Curr Opin Cell Biol 62, 9–16 (2020).

4. Gillies, T.E. & Cabernard, C. Cell division orientation in animals. Curr Biol 21, R599–609 (2011).

5. Lu, M.S. & Johnston, C.A. Molecular pathways regulating mitotic spindle orientation in animal cells. Development 140, 1843–1856 (2013).

6. di Pietro, F., Echard, A. & Morin, X. Regulation of mitotic spindle orientation: an integrated view. EMBO reports 17, 1106–1130 (2016).

7. Seldin, L. & Macara, I. Epithelial spindle orientation diversities and uncertainties: recent developments and lingering questions. F1000Research 6, 984–984 (2017).

8. Bergstralh, D.T., Dawney, N.S. & St Johnston, D. Spindle orientation: a question of complex positioning. Development 144, 1137–1145 (2017).

9. Dewey, E.B., Taylor, D.T. & Johnston, C.A. Cell Fate Decision Making through Oriented Cell Division. J Dev Biol 3, 129–157 (2015).

10. Bergstralh, D.T. & St Johnston, D. Spindle orientation: what if it goes wrong? Semin Cell Dev Biol 34, 140–145 (2014).

11. Niwayama, R. et al. A Tug-of-War between Cell Shape and Polarity Controls Division Orientation to Ensure Robust Patterning in the Mouse Blastocyst. Developmental Cell 51, 564–574.e566 (2019).

12. Li, J., Cheng, L. & Jiang, H. Cell shape and intercellular adhesion regulate mitotic spindle orientation. Mol Biol Cell 30, 2458–2468 (2019).

13. Petridou, N.I. & Skourides, P.A. A ligand-independent integrin beta1 mechanosensory complex guides spindle orientation. Nature communications 7, 10899 (2016).

14. Dimitracopoulos, A. et al. Mechanochemical Crosstalk Produces Cell-Intrinsic Patterning of the Cortex to Orient the Mitotic Spindle. Curr Biol 30, 3687–3696 e3684 (2020).

15. Li, Y. & Burridge, K. Cell-Cycle-Dependent Regulation of Cell Adhesions: Adhering to the Schedule: Three papers reveal unexpected properties of adhesion structures as cells progress through the cell cycle. Bioessays 41, e1800165 (2019).

16. Lesman, A., Notbohm, J., Tirrell, D.A. & Ravichandran, G. Contractile forces regulate cell division in three-dimensional environments. J Cell Biol 205, 155–162 (2014).

17. Matsumura, S. et al. Interphase adhesion geometry is transmitted to an internal regulator for spindle orientation via caveolin-1. Nature communications 7, ncomms11858 (2016).

18. Rizzelli, F., Malabarba, M.G., Sigismund, S. & Mapelli, M. The crosstalk between microtubules, actin and membranes shapes cell division. Open biology 10, 190314 (2020).

19. Mitchison, T.J. Actin based motility on retraction fibers in mitotic PtK2 cells. Cell Motil Cytoskeleton 22, 135–151 (1992).

20. Charnley, M., Anderegg, F., Holtackers, R., Textor, M. & Meraldi, P. Effect of Cell Shape and Dimensionality on Spindle Orientation and Mitotic Timing. PLoS One 8, e66918 (2013).

21. Thery, M. & Bornens, M. Cell shape and cell division. Curr Opin Cell Biol 18, 648–657 (2006).

22. Nestor-Bergmann, A. et al. Decoupling the Roles of Cell Shape and Mechanical Stress in Orienting and Cueing Epithelial Mitosis. Cell reports 26, 2088–2100 e2084 (2019).

23. Walma, D.A.C. & Yamada, K.M. The extracellular matrix in development. Development 147, dev175596 (2020).

24. Anastasiou, O., Hadjisavva, R. & Skourides, P.A. Mitotic cell responses to substrate topological cues are independent of the molecular nature of adhesion. Science Signaling 13, eaax9940 (2020).

25. Fink, J. et al. External forces control mitotic spindle positioning. Nat Cell Biol 13, 771–778 (2011).

26. Thery, M. et al. The extracellular matrix guides the orientation of the cell division axis. Nat Cell Biol 7, 947–953 (2005).

27. Finegan, T.M. & Bergstralh, D.T. Division orientation: disentangling shape and mechanical forces. Cell Cycle 18, 1187–1198 (2019).

28. Lam, M.S.Y. et al. Isotropic myosin-generated tissue tension is required for the dynamic orientation of the mitotic spindle. Mol Biol Cell 31, 1370–1379 (2020).

29. Nestor-Bergmann, A., Goddard, G. & Woolner, S. Force and the spindle: mechanical cues in mitotic spindle orientation. Semin Cell Dev Biol 34, 133–139 (2014).

30. Kotak, S. & Gonczy, P. Mechanisms of spindle positioning: cortical force generators in the limelight. Curr Opin Cell Biol (2013).

31. Gibson, M.C., Patel, A.B., Nagpal, R. & Perrimon, N. The emergence of geometric order in proliferating metazoan epithelia. Nature 442, 1038–1041 (2006).

32. Pinheiro, D. & Bellaiche, Y. Mechanical Force-Driven Adherens Junction Remodeling and Epithelial Dynamics. Dev Cell 47, 3–19 (2018).

33. Guillot, C. & Lecuit, T. Mechanics of epithelial tissue homeostasis and morphogenesis. Science 340, 1185–1189 (2013).

34. Cramer, L.P. & Mitchison, T.J. Myosin is involved in postmitotic cell spreading. J Cell Biol 131, 179–189 (1995).

35. Higashi, T. & Miller, A.L. Tricellular junctions: how to build junctions at the TRICkiest points of epithelial cells. Mol Biol Cell 28, 2023–2034 (2017).

36. Lecuit, T. & Yap, A.S. E-cadherin junctions as active mechanical integrators in tissue dynamics. Nat Cell Biol 17, 533–539 (2015).

37. Pannekoek, W.J., de Rooij, J. & Gloerich, M. Force transduction by cadherin adhesions in morphogenesis. F1000Research 8 (2019).

38. Shimoyama, Y. et al. Cadherin cell-adhesion molecules in human epithelial tissues and carcinomas. Cancer Res 49, 2128–2133 (1989).

39. Tomlinson, J.S., Alpaugh, M.L. & Barsky, S.H. An intact overexpressed E-cadherin/alpha,beta-catenin axis characterizes the lymphovascular emboli of inflammatory breast carcinoma. Cancer Res 61, 5231–5241 (2001).

40. Wang, X. et al. E-cadherin bridges cell polarity and spindle orientation to ensure prostate epithelial integrity and prevent carcinogenesis in vivo. PLOS Genetics 14, e1007609 (2018).

41. Hart, K.C. et al. E-cadherin and LGN align epithelial cell divisions with tissue tension independently of cell shape. Proc Natl Acad Sci U S A 114, E5845–E5853 (2017).

42. Gloerich, M., Bianchini, J.M., Siemers, K.A., Cohen, D.J. & Nelson, W.J. Cell division orientation is coupled to cell-cell adhesion by the E-cadherin/LGN complex. Nature communications 8, 13996 (2017).

43. den Elzen, N., Buttery, C.V., Maddugoda, M.P., Ren, G. & Yap, A.S. Cadherin adhesion receptors orient the mitotic spindle during symmetric cell division in mammalian epithelia. Mol Biol Cell 20, 3740–3750 (2009).

44. Kotak, S. Mechanisms of Spindle Positioning: Lessons from Worms and Mammalian Cells. Biomolecules 9 (2019).

45. Machicoane, M. et al. SLK-dependent activation of ERMs controls LGN-NuMA localization and spindle orientation. J Cell Biol 205, 791–799 (2014).

46. Yamashita, Y.M., Jones, D.L. & Fuller, M.T. Orientation of asymmetric stem cell division by the APC tumor suppressor and centrosome. Science 301, 1547–1550 (2003).

47. di Pietro, F., Echard, A. & Morin, X. Regulation of mitotic spindle orientation: an integrated view. EMBO Rep 17, 1106–1130 (2016).

48. Taneja, N. et al. Focal adhesions control cleavage furrow shape and spindle tilt during mitosis. Scientific reports 6, 29846 (2016).

49. Cavey, M., Rauzi, M., Lenne, P.F. & Lecuit, T. A two-tiered mechanism for stabilization and immobilization of E-cadherin. Nature 453, 751–756 (2008).

50. Godde, N.J., et al. Scribble modulates the MAPK/Fra1 pathway to disrupt luminal and ductal integrity and suppress tumour formation in the mammary gland. PLoS Genet 10, e1004323 (2014).

51. Nakajima, Y.I., Meyer, E.J., Kroesen, A., McKinney, S.A. & Gibson, M.C. Epithelial junctions maintain tissue architecture by directing planar spindle orientation. Nature (2013).

52. Bonello, T.T. & Peifer, M. Scribble: A master scaffold in polarity, adhesion, synaptogenesis, and proliferation. J Cell Biol 218, 742–756 (2019).

53. Qin, Y., Capaldo, C., Gumbiner, B.M. & Macara, I.G. The mammalian Scribble polarity protein regulates epithelial cell adhesion and migration through E-cadherin. J Cell Biol 171, 1061–1071 (2005).

54. Nakajima, Y.-i. et al. Junctional tumor suppressors interact with 14-3-3 proteins to control planar spindle alignment. Journal of Cell Biology 218, 1824–1838 (2019).

55. Navarro, C. et al. Junctional recruitment of mammalian Scribble relies on E-cadherin engagement. Oncogene 24, 4330–4339 (2005).

56. Allam, A.H., Charnley, M. & Russell, S.M. Context-Specific Mechanisms of Cell Polarity Regulation. J Mol Biol 430, 3457–3471 (2018).

57. Zigman, M., Trinh le, A., Fraser, S.E. & Moens, C.B. Zebrafish neural tube morphogenesis requires Scribble-dependent oriented cell divisions. Curr Biol 21, 79–86 (2011).

58. Bell, G.P., Fletcher, G.C., Brain, R. & Thompson, B.J. Aurora Kinases Phosphorylate Lgl to Induce Mitotic Spindle Orientation in Drosophila Epithelia. Curr Biol 25, 61–68 (2015).

59. Carvalho, C.A., Moreira, S., Ventura, G., Sunkel, C.E. & Morais-de-Sa, E. Aurora A triggers Lgl cortical release during symmetric division to control planar spindle orientation. Curr Biol 25, 53–60 (2015).

60. Porter, A.P., White, G.R.M., Mack, N.A. & Malliri, A. The interaction between CASK and the tumour suppressor Dlg1 regulates mitotic spindle orientation in mammalian epithelia. Journal of Cell Science, jcs.230086 (2019).

61. Thery, M., Jimenez-Dalmaroni, A., Racine, V., Bornens, M. & Julicher, F. Experimental and theoretical study of mitotic spindle orientation. Nature 447, 493–496 (2007).

62. Kiyomitsu, T. The cortical force-generating machinery: how cortical spindle-pulling forces are generated. Current Opinion in Cell Biology 60, 1–8 (2019).

63. Du, Q. & Macara, I.G. Mammalian Pins is a conformational switch that links NuMA to heterotrimeric G proteins. Cell 119, 503–516 (2004).

64. Zheng, Z., Wan, Q., Meixiong, G. & Du, Q. Cell cycle-regulated membrane binding of NuMA contributes to efficient anaphase chromosome separation. Mol Biol Cell 25, 606–619 (2014).

65. Lock, J.G. & Stow, J.L. Rab11 in recycling endosomes regulates the sorting and basolateral transport of E-cadherin. Mol Biol Cell 16, 1744–1755 (2005).

66. Johnston, C.A., Hirono, K., Prehoda, K.E. & Doe, C.Q. Identification of an Aurora-A/PinsLINKER/Dlg spindle orientation pathway using induced cell polarity in S2 cells. Cell 138, 1150–1163 (2009).

67. Bergstralh, D.T., Lovegrove, H.E. & St Johnston, D. Discs large links spindle orientation to apical-basal polarity in Drosophila epithelia. Curr Biol 23, 1707–1712 (2013).

68. Saadaoui, M. et al. Dlg1 controls planar spindle orientation in the neuroepithelium through direct interaction with LGN. J Cell Biol 206, 707–717 (2014).

69. Rathbun, L.I. et al. Cytokinetic bridge triggers de novo lumen formation in vivo. Nature communications 11, 1269 (2020).

70. Fededa, J.P. & Gerlich, D.W. Molecular control of animal cell cytokinesis. Nat Cell Biol 14, 440–447 (2012).

71. Papalazarou, V. & Machesky, L.M. The cell pushes back: The Arp2/3 complex is a key orchestrator of cellular responses to environmental forces. Current Opinion in Cell Biology 68, 37–44 (2021).

72. Herszterg, S., Leibfried, A., Bosveld, F., Martin, C. & Bellaiche, Y. Interplay between the dividing cell and its neighbors regulates adherens junction formation during cytokinesis in epithelial tissue. Dev Cell 24, 256–270 (2013).

73. Li, J.X.H., Tang, V.W. & Brieher, W.M. Actin protrusions push at apical junctions to maintain E-cadherin adhesion. Proc Natl Acad Sci U S A 117, 432–438 (2020).

74. Kwon, M., Bagonis, M., Danuser, G. & Pellman, D. Direct Microtubule-Binding by Myosin-10 Orients Centrosomes toward Retraction Fibers and Subcortical Actin Clouds. Developmental cell 34, 323–337 (2015).

75. Gallop, J.L. Filopodia and their links with membrane traffic and cell adhesion. Semin Cell Dev Biol 102, 81–89 (2020).

76. He, K., Sakai, T., Tsukasaki, Y., Watanabe, T.M. & Ikebe, M. Myosin X is recruited to nascent focal adhesions at the leading edge and induces multi-cycle filopodial elongation. Scientific reports 7, 13685 (2017).

77. Cramer, L. & Mitchison, T.J. Moving and stationary actin filaments are involved in spreading of postmitotic PtK2 cells. J Cell Biol 122, 833–843 (1993).

78. Li, Y. et al. Discs large 1 controls daughter-cell polarity after cytokinesis in vertebrate morphogenesis. Proc Natl Acad Sci U S A 115, E10859–E10868 (2018).

79. Dix, C.L. et al. The Role of Mitotic Cell-Substrate Adhesion Re-modeling in Animal Cell Division. Dev Cell 45, 132–145 e133 (2018).

80. Lock, J.G. et al. Reticular adhesions are a distinct class of cell-matrix adhesions that mediate attachment during mitosis. Nat Cell Biol 20, 1290–1302 (2018).

81. Charnley, M., Kroschewski, R. & Textor, M. The study of polarisation in single cells using model cell membranes. Integr Biol (Camb) 4, 1059–1071 (2012).

82. Thompson, C.J. et al. Cadherin clusters stabilized by a combination of specific and nonspecific cis-interactions. eLife 9, e59035 (2020).

83. Thompson, C.J., Vu, V.H., Leckband, D.E. & Schwartz, D.K. Cadherin cis and trans interactions are mutually cooperative. Proceedings of the National Academy of Sciences 118, e2019845118 (2021).

84. Thompson, C.J., Vu, V.H., Leckband, D.E. & Schwartz, D.K. Cadherin Extracellular Domain Clustering in the Absence of Trans-Interactions. The journal of physical chemistry letters 10, 4528–4534 (2019).

85. Discher, D.E., Janmey, P. & Wang, Y.L. Tissue cells feel and respond to the stiffness of their substrate. Science 310, 1139–1143 (2005).

86. Uroz, M. et al. Regulation of cell cycle progression by cell-cell and cell-matrix forces. Nat Cell Biol 20, 646–654 (2018).

87. Osswald, M. & Morais-de-Sa, E. Dealing with apical-basal polarity and intercellular junctions: a multidimensional challenge for epithelial cell division. Curr Opin Cell Biol 60, 75–83 (2019).

88. Le Bras, S. & Le Borgne, R. Epithelial cell division - multiplying without losing touch. J Cell Sci 127, 5127–5137 (2014).

89. Sharifkhodaei, Z., Gilbert, M.M. & Auld, V.J. Scribble and Discs Large mediate tricellular junction formation. Development 146, dev174763 (2019).

90. Bosveld, F. & Bellaiche, Y. Tricellular junctions. Curr Biol 30, R249–R251 (2020).

91. Wang, Z., Bosveld, F. & Bellaiche, Y. Tricellular junction proteins promote disentanglement of daughter and neighbour cells during epithelial cytokinesis. J Cell Sci 131 (2018).

92. Trylinski, M. & Schweisguth, F. Activation of Arp2/3 by WASp Is Essential for the Endocytosis of Delta Only during Cytokinesis in Drosophila. Cell reports 28, 1–10.e13 (2019).

93. Dow, L.E. et al. The tumour-suppressor Scribble dictates cell polarity during directed epithelial migration: regulation of Rho GTPase recruitment to the leading edge. Oncogene 26, 2272–2282 (2007).

94. Humbert, P.O., Dow, L.E. & Russell, S.M. The Scribble and Par complexes in polarity and migration: friends or foes? Trends Cell Biol 16, 622–630 (2006).

95. Ludford-Menting, M.J. et al. A network of PDZ-containing proteins regulates T cell polarity and morphology during migration and immunological synapse formation. Immunity 22, 737–748 (2005).

96. Osmani, N., Vitale, N., Borg, J.P. & Etienne-Manneville, S. Scrib controls Cdc42 localization and activity to promote cell polarization during astrocyte migration. Curr Biol 16, 2395–2405 (2006).

