## Supplementary material for "A Scribble-E-cadherin complex controls daughter cell patterning by multiple mechanisms": Methods

**Cell line and cell culture**

MCF10A cells were cultured in DMEM:F12 (Dulbecco's Modified Eagle Medium: F12) media, supplemented with 5% (v/v) horse serum (Gibco), 10 μg/ml insulin (Novo Nordisk) , 0.5 μg/ml hydrocortisone (Sigma-Aldrich), 20 ng/ml human epidermal growth factor (Sigma-Aldrich), 100 ng/ml cholera toxin (List Biological Labs), 100 ng/ml penicillin / streptomycin, 2 mM glutamine, and maintained at 37°C in 5% CO_2_. Stable MCF10A cell lines that expressed short hairpin RNA against human Scribble (#7) and the control were created and analysed as described in ^1^. Scribble depletion was re-confirmed by flow cytometry (Supplementary Figure.2a). Briefly, the cells were fixed by IC Fixation Buffer (eBioscience) according to manufacturer’s protocol, followed by incubation with anti-Scribble antibody (sc-11049, Santa Cruz Biotechnology), and then labelling with Alexa Fluor 647 conjugated antibody (ab150131, Abcam), and examined by flow cytometry (BD FACSAria III, BD).

HeLa cells were cultured in DMEM media (Gibco), supplemented with 10% (v/v) fetal bovine serum, 2 mM glutamine and 100 ng/ml penicillin / streptomycin, and maintained at 37°C in 5% CO_2_.

**si-RNA silencing**

Scribble of MCF10A cells and HeLa cells was depleted by siGENOME SMARTpool si-RNA against human Scribble. si-RNA was carried by DharmaFECT 3 according to the manufacturer’s protocol, and was incubated with cells in serum-free cell culture media for transfection. We applied two rounds of si-RNA silencing to increase knockdown efficiency. Briefly, the first round of si-RNA silencing (20 nM of si-RNA) was applied on a 70% confluent monolayer cultured in a well of a 12-well culture plate. After an incubation with si-RNA containing media for 16 hours, the cells were refreshed with regular culture media, and incubated for further 8 hours. We plated these si-RNA-transfected cells on glass-bottomed 8-well chamber (ibidi) for the purpose of subsequent cell imaging, and grew the cells for around 12 hours. A second round of 8-hour si-RNA transfection (10 nM of si-RNA) was applied to these cells, and a subsequent 16-hour incubation under regular culture media. Cells were then fixed for further analysis.

**E-cadherin blocking treatment**

Mouse monoclonal anti-E-cadherin antibody HECD-1 (ab1416, Abcam) was used as to block the function of surface E-cadherin. For the fixed-cell imaging, the sparsely plated cell (2× 103 cells/cm^2^) was cultured in the culture media containing 5 μg/ml antibody HECD-1 for 1.5 hour before fixation. For the live-cell imaging, the sparsely plated cell (2× 10^3^ cells/cm^2^) was cultured in the HECD-1 containing culture media (5 μg/ml) for one hour prior the time-lapse acquisition, followed with image acquisition for around 2 hours.

**Drug treatment**

Actin depolymerisor Cytochalasin D (Sigma-Aldrich) was used at 5 μM, Myosin II inhibitor Blebbistatin (Sigma-Aldrich) was used at 2.5 μM, and Arp2/3 inhibitor CK-869 (Sigma-Aldrich) was used at 50 μM. For fixed imaging, the inhibitors were applied for 15 minutes under regular culture condition before fixation. For live-cell imaging, Arp2/3 inhibitor CK-869 was used at 0.1 μM for 2 hours during image acquisition.

**Immunofluorescence staining and confocal microscopy for imaging of fixed cells**

MCF10A cells were grown on glass-bottomed 8-well chambers (ibidi), and fixed with 4% formaldehyde for 8 minutes at room temperature followed by permeabilization with 0.1% Triton X-100 for 5 minutes. The samples were then incubated with primary antibody overnight at 4°C as described in Supplementary table 1. Next, the samples were washed with PBS, and labelled with fluorochrome-conjugated secondary antibodies (Supplementary table 1), DAPI (Thermo Fisher Scientific), and Phalloidin (Abcam). The samples were examined using a FV3000 confocal microscope (Olympus) and 60X lens (1.30 NA, UPLSAPO, Olympus). To acquire images of the whole cell, scanning of multiple sections with a z distance of 0.5 μm was applied. These sections were either merged or converted to different orthogonal projections using Image J, and were also used in the analysis of spindle orientation. To acquire images of subcellular structures such as retraction fibres or filopodia, scanning of multiple sections with a z distance of 0.2 μm was applied. The images were processed with maximum intensity projection of three continuous stages for subcellular structures such as retraction fibres, filopodia and nascent junctions using Image J. An ImageJ plugin ‘3D viewer’ was used to reconstruct the three-dimensional images.

**Time-lapse microscopy for live cells imaging**

For live-cell imaging, MCF10A cells were sparsely plated at around 2× 10^3^ cells/cm^2^ and cultured for 1.5 days. Before image acquisition, the cells were incubated in cell culture media containing 250 nM SiR-DNA (Spirochrome) to label DNA for 30 minutes, which was then replaced with regular cell culture media. The process of mitosis and cell division were imaged using an Olympus FV3000 confocal microscope (Olympus) and 20X lens (0.75 NA, UPLSAPO, Olympus) in a chamber maintained in 37°C and with 5% CO_2_. Mitotic cells were identified by the rounded cell morphology and the bar shape of condensed chromosomes, and were recorded every 3 minutes for more than 30 minutes after anaphase onset. The z scanning, with 1 µm per z stage, was applied to cover the whole volume of the cell. The images were processed with average intensity projection for both bright field channel and the channel for SiR-DNA by Image J. To quantify the time taken by each daughter to re-adhere to the substrate, the time frame showing each daughter cell flattening was identified as the time of re-adherence, and the time of entry into anaphase onset was subtracted. To quality the distance between daughters’ nucleus labelled by SiR-DNA, we measured the shortest distance between two segregated parts of DNA.

**Data collection and analysis**

Confocal images were collected using a Olympus FV3000 confocal microscope and FV31S-SW Viewer software, and were processed using ImageJ (version 1.52p).

Flow cytometry data were collected using a BD FACSAria III and software BD FACSDiVa (version 6.1.3), and were processed using FlowJo (version X).

ImageJ (version 1.52p) was used for quantification on confocal images; Python (version 3.8.1) was used for statistics; FlowJo (version X) was used for analysing flow cytometry data; Microsoft Excel 2016 was used for data organisation and t-test.

The following algorithms were used in Python for analysis and plotting: matplotlib (version 3.1.3), pandas (version 1.0.1), seaborn (version 0.10.0), numpy (version 1.18.1), and plotly (version 4.14.3).

The bar plot was shown as median ± one quartile. We used t-test to determine the significance between two distributions of variables using the built-in formula in Microsoft Office Excel. The results were derived from at least three independent experiments.

1.	Dow, L.E. *et al.* The tumour-suppressor Scribble dictates cell polarity during directed epithelial migration: regulation of Rho GTPase recruitment to the leading edge. *Oncogene* **26**, 2272-2282 (2007).
