## Supplementary material for "A Scribble-E-cadherin complex controls daughter cell patterning by multiple mechanisms": Methods: Antibody table

| Antibody | Supplier | Catlog | Usage (titre) | |
|---|---|---|---|---|
| Integrin β1 | Abcam | ab30394 | Immunofluorescence | 1:300 |
| E-cadherin | Abcam | ab1416 | Immunofluorescence | 1:100 |
| E-cadherin | Cell Signaling Technology | 3195 | Immunofluorescence | 1:250 |
| α-Tubulin | Rockland Immunochemicals | 200-301-880 | Immunofluorescence | 1:1000 |
| α-Tubulin | Rockland Immunochemicals | 600-401-880 | Immunofluorescence | 1:1000 |
| Scribble | Santa Cruz Biotechnology | sc-11049 | Immunofluorescence | 1:250 |
| Scribble | Upstate | 07-643 | Immunofluorescence | 1:500 |
| NuMA | Abcam | ab109262 | Immunofluorescence | 1:500 |
| Paxillin | Millipore | 05-417 | Immunofluorescence | 1:400 |
| Arp2 | Abcam | ab49674 | Immunofluorescence | 1:500 |
| Ezrin | BD Biosciences | 610602 | Immunofluorescence | 1:800 |
| Myosin X | Novus Biologicals | NBP1-87748 | Immunofluorescence | 1:500 |
| β-Catenin | BD Biosciences | 610153 | Immunofluorescence | 1:500 |
| β-PIX | Cell Signaling Technology | 4515 | Immunofluorescence | 1:400 |
| β-PIX | Millipore | 07-1450-I | Immunofluorescence | 1:500 |
| Myosin IIb | Cell Signaling Technology | 8824 | Immunofluorescence | 1:500 |
